# Acute lysergic acid diethylamide induces a time-dependent shift toward hippocampal control of default mode network reorganization

**DOI:** 10.64898/2026.09.03.748807

**Authors:** Fahd François Hilal, Rayane Benkeddada, Jerome Jeanblanc, Marion Sourty, Sidy Fall, Rachel Utama, Sima Soltanpour, Praveen Kulkarni, Mickael Naassila, Sami Ben Hamida, Md Taufiq Nasseef

**Author notes:** Correspondence to: Md Taufiq Nasseef, Department of Mathematics, College of Science and Humanity Studies, Prince Sattam Bin Abdulaziz University, Al Kharj, Riyadh, Saudi Arabia, work; Correspondence to: Sami Ben Hamida, Mickael Naassila, Alcohol and Pharmacodependence Research Group, UMR INSERM U1247, University of Picardie Jules Verne, Amiens, France, work. Co-first authors, these authors contributed equally to this work (Fahd François Hilal, Rayane Benkeddada). Co-last authors, these authors contributed equally to this work (Mickael Naassila, Sami Ben Hamida, Md Taufiq Nasseef).

## Abstract

**Background:** Classical psychedelics induce profound changes in brain function, yet the temporal organization of these effects remains incompletely understood.

**Methods:** We investigated the effects of acute lysergic acid diethylamide (LSD) on large-scale brain connectivity in nine male Long–Evans rats using resting-state functional magnetic resonance imaging. Following a 15-min baseline acquisition, rats received LSD (500 µg/kg, i.p.), and imaging continued during two consecutive 15-min post-injection periods. Network reorganization was assessed using independent component analysis (ICA), seed-to-voxel mapping, stationary seed-to-seed connectivity, dynamic graph-based metrics, and dynamic causal modeling.

**Results:** LSD produced widespread but regionally heterogeneous changes across default mode network– related, cortical, striatal, thalamic, sensory, and limbic systems. During the early post-injection window, ICA revealed a predominance of increased functional coupling, while dynamic analyses showed prominent fluctuations in hippocampal coupling strength and medial frontal network centrality. Dynamic causal modeling identified a false discovery rate–corrected reduction in retrosplenial-to-infralimbic influence, accompanied by a broader descriptive pattern of reduced posterior-and hippocampal-to-frontal coupling. During the later window, ICA showed a relative shift toward connectivity decreases, while complementary analyses revealed more selective hippocampal and parahippocampal involvement.

**Conclusions:** Across complementary analyses, acute LSD induced a temporally structured reorganization of brain hierarchy, characterized by an early weakening of directed cortical interactions followed by a later shift toward hippocampal-centered network control. These findings identify a dynamic shift in effective network organization and provide a mechanistic framework for psychedelic-induced default mode network reconfiguration.

## Introduction

Over the past two decades, scientific interest in psychedelic compounds has resurged, driven by renewed clinical promise and by the need to better define the neurobiological mechanisms underlying their effects (1). Lysergic acid diethylamide (LSD), a prototypical classical psychedelic, produces profound and reliable alterations in perception, cognition, and self-related processing (2). Although these effects are primarily attributable to cortical 5-HT2A receptor activation (3) and downstream glutamatergic signaling (4), they extend beyond local receptor-level mechanisms to coordinated changes in distributed brain systems (1).

Human neuroimaging studies consistently report that acute classical psychedelics alter large-scale functional organization, with reduced resting-state network segregation and increased communication between normally distinct systems (3,5,6). Recent large-scale evidence identifies increased transmodal–unimodal/sensorimotor coupling, selective rather than global within-network reductions, and reproducible subcortical involvement extending beyond the thalamus to the caudate and putamen as core features of psychedelic-induced network reorganization (7). At the network level, earlier psilocybin studies reported reduced activity and connectivity in medial default mode network (DMN) hubs, particularly the medial prefrontal cortex (mPFC) and posterior cingulate cortex (PCC), together with increased communication between normally segregated networks (6,8,9). Similarly, LSD induces large-scale connectivity changes consistent with network reorganization (3,5,10,11). Most early studies, however, relied on stationary resting-state functional connectivity averaged over long time windows. More recent dynamic analyses have demonstrated that LSD-induced changes in network integration and segregation are non-uniform over time (12). Because functional connectivity captures statistical dependencies but not directionality, effective connectivity can estimate directed influences within defined circuits and provide a more mechanistic account of psychedelic-induced network reorganization (13). Using spectral dynamic causal modelling, LSD-induced alterations in directed connectivity within cortico-striato-thalamo-cortical pathways were identified (14). More recently, whole-brain regression DCM revealed predominantly stronger interregional effective connectivity under LSD, together with regionally heterogeneous changes in inhibitory self-connections and local gain (15).

Psychedelics have been shown to increase dynamical complexity and expand the repertoire of network configurations explored by the brain (12,16). Although medial cortical and hippocampal regions are consistently implicated, how their contributions evolve across the acute psychedelic state remains incompletely resolved (12,16,17). Preclinical neuroimaging provides an experimentally controlled framework through precise dosing and repeated measurements across defined post-drug phases. In rodents, psilocybin, psilocin, and LSD induce widespread alterations in brain activity and resting-state connectivity across cortical, striatal, thalamic, and limbic systems, broadly paralleling systems-level effects observed in humans (18–21). However, the evolution of these changes across successive phases of the acute LSD response remains poorly characterized.

In the present study, we investigated the onset and early evolution of LSD-induced large-scale brain connectivity reorganization during the initial 30-min post-injection period in rats, using a multimodal framework combining independent component analysis, seed-to-seed connectivity, dynamic graph-centrality metrics, effective connectivity, and seed-to-voxel mapping. We hypothesized that LSD would induce a temporally structured reconfiguration of large-scale network architecture involving default-mode, hippocampal, thalamic, limbic and sensory-associative systems rather than a stationary disruption of connectivity. By resolving this reorganization across consecutive post-injection periods and complementary connectivity dimensions, we tested whether apparently distinct network signatures represent successive phases of a common evolving process.

## Methods and Materials

### Animals

Nine male Long–Evans rats (9 weeks old; 246–332 g at MRI acquisition; Janvier Labs, Le Genest-Saint-Isle, France) were individually housed with ad libitum food and water. Procedures complied with Directive 2010/63/EU and were approved by the local ethics committee (CREMEAP; APAFIS 26615); the study is reported in accordance with ARRIVE guidelines. Further details are provided in Supplemental Methods.

### Drugs

LSD tartrate (LGC Standards; CAS No. 32426-57-6) was dissolved in sterile saline (0.9% NaCl) containing 1% DMSO and administered intraperitoneally (i.p.) at 500 µg/kg, expressed as free-base equivalent, at an injection volume of 1 mL/kg.

### Image acquisition

The individual rat was the experimental unit (n = 9), with repeated measurements across the three imaging periods. No formal a priori sample-size calculation was performed; sample size was based on feasibility, prior rodent rs-fMRI experience, and comparable studies (18,19). A within-subject repeated-measures design used each animal as its own baseline control; no time-matched vehicle group was included. Rats were anesthetized with isoflurane (5% induction; 1.5–1.8% maintenance) and their head secured. Temperature, respiration, oxygen saturation, and heart rate were continuously monitored and remained within physiological ranges (Supplementary Figure 1).

MRI was performed on a 7T Bruker BioSpec 70/20 USR scanner. Coronal BOLD images were acquired using SE-EPI (TR/TE = 2000/16 ms; flip angle = 90°; 450 volumes; 34 slices; 0.8-mm thickness). Each session included anatomical imaging, a 15-min baseline run, a 90-s pause for remote i.p. LSD administration without repositioning, and two consecutive 15-min post-injection runs. Isoflurane was increased briefly to 2% prior to injection, then reduced to 1.5–1.8% before acquisition resumed. Further acquisition and physiological-monitoring details are provided in Supplemental Methods; Figure 1 summarizes the design.

**Figure 1.**
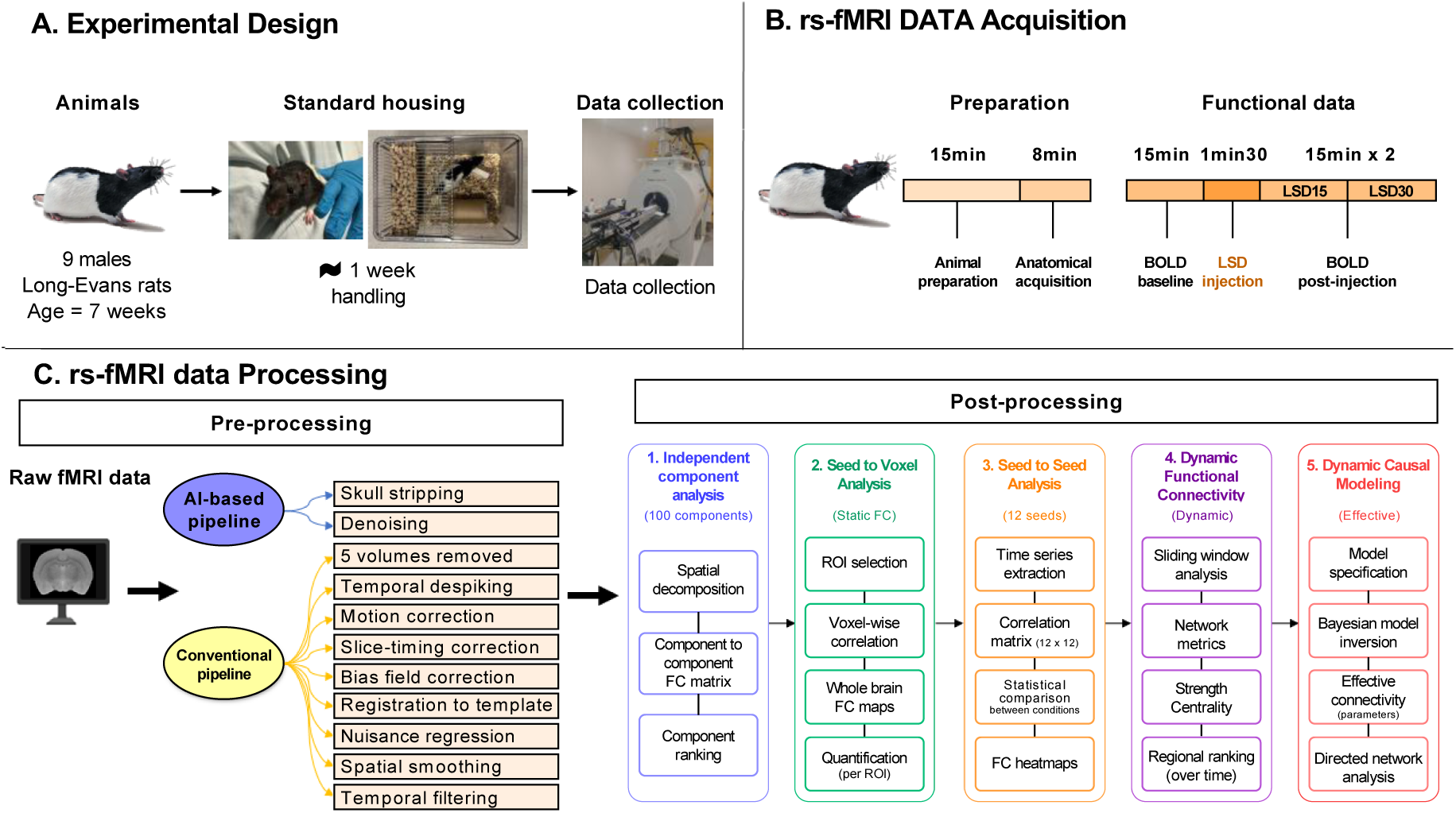
Experimental design, rs-fMRI acquisition, preprocessing, and analytical workflow. A) Experimental design. Nine male Long–Evans rats were included in the study. Animals arrived at 7 weeks of age and were maintained under standard housing conditions with handling before imaging. Resting-state fMRI data were acquired at 9 weeks of age on a 7 T Bruker BioSpec system under controlled isoflurane anesthesia. B) rs-fMRI acquisition protocol. Animals were prepared under isoflurane anesthesia, followed by anatomical and resting-state functional imaging. Each rat underwent a 15-min baseline resting-state scan, followed by intraperitoneal administration of LSD tartrate (500 µg/kg free-base equivalent; 1 mL/kg; 0.9% NaCl + 1% DMSO) using a remote injection system without repositioning the animal. Functional imaging resumed 90 s after injection and continued for two consecutive 15-min post-injection runs, hereafter referred to as LSD-15 and LSD-30. All animals completed the full protocol and were included in the analyses (n = 9). Physiological parameters were continuously monitored throughout acquisition. C) rs-fMRI preprocessing and analysis workflow. Raw functional data were converted to NIfTI format and processed using an automated pipeline combining deep-learning and conventional preprocessing steps. Preprocessing included skull stripping, denoising, removal of the first five volumes, despiking, motion correction, slice-timing correction, bias-field correction, registration to a 174-region 3D MRI Rat Brain Atlas, nuisance regression, spatial smoothing, and temporal band-pass filtering. Postprocessing analyses included independent component analysis (ICA), seed-to-voxel functional connectivity (FC), seed-to-seed stationary FC, dynamic FC (DFC), and dynamic causal modeling (DCM). These complementary analyses were used to assess LSD-associated changes in large-scale functional organization, regional connectivity, temporal network dynamics, and directed effective connectivity.

A priori exclusion criteria included unsuccessful LSD administration, excessive motion, physiological instability, failed registration, or poor BOLD data quality. No dataset met these criteria; all nine rats were retained in every analysis. The order in which rats were scanned was randomized. Investigators were aware of the imaging period during data acquisition and analysis. Data analyses were conducted using predefined standardized pipelines.

### Image preprocessing

Raw Bruker fMRI data were converted to NIfTI format and processed using an automated preprocessing pipeline. Skull stripping was performed using a Smart Swin Transformer combined with a Dense U-Net architecture (22), followed by denoising using a 3D U-WGAN (23). After discarding the first five volumes, data underwent despiking, motion and slice-timing correction, bias-field correction, and registration to an in-house 174-region 3D MRI Rat Brain Atlas (© Ekam Solutions, Boston, MA, USA). Nuisance regression included motion parameters and mean ventricular and white-matter signals (22,24,25). Data were spatially smoothed (0.4-mm FWHM) and band-pass filtered (0.01–0.1 Hz); full procedures are provided in Supplemental Methods.

### Post-processing

Resting-state fMRI data were examined using complementary approaches comprising high-dimensional independent component analysis (ICA; 100 components), seed-to-seed stationary and dynamic functional connectivity, dynamic causal modeling, and whole-brain seed-to-voxel analysis (26–28). Further analytical details are provided in Supplemental Methods.

### Independent Component Analysis

Group ICA was performed in GIFT using all rats and sessions (baseline, LSD 15 min, and LSD 30 min) as a single dataset (26–28). One hundred components were estimated, with stability assessed by ICASSO (20 repetitions). Correlations between anatomically annotated component time series were Fisher Z-transformed. Within-session component correlations were tested against zero (one-sample t tests; t > 4, p < 0.01; n = 9). Baseline was compared separately with LSD 15 min and LSD 30 min using pairwise two-tailed t-tests (t > 2; p = 0.05), with FDR correction across component pairs. Component importance was ranked by the number of significant FC changes involving each component (27).

### Seed-to-voxel analysis

Seed-to-voxel FC was assessed for six predefined seeds: anterior cingulate cortex (ACC), hippocampus (HPC), orbitofrontal cortex (OFC), parahippocampal region (paraHPC), medial prefrontal cortex (mPFC), and retrosplenial cortex (RSC). For each rat and session, mean seed BOLD time series were correlated voxel-wise with the rest of the brain and Fisher Z-transformed. Within-session positive connectivity was assessed using one-sample t tests (t > 4, p < 0.01; n = 9). Baseline was compared separately with LSD 15 min and LSD 30 min using directional t tests (p < 0.05), with cluster-level familywise error correction (pFWE < 0.05). Mean Z-values were extracted from selected anatomical regions, and seed-target effects were additionally quantified descriptively by the proportion of significant voxels for ranking.

### Seed-to-seed stationary FC analysis

Seed-to-seed stationary FC was assessed across 12 predefined regions spanning DMN-related, sensory-associative, and serotonergic systems. The rat DMN-related set comprised OFC, ACC, prelimbic cortex (PL), infralimbic cortex (IL), auditory cortex (AUD), HPC, temporal association cortex (TempA), RSC, parietal association cortex (Par), and visual cortex (VIS), based on established rodent DMN organization and its application in psychedelic pharmaco-fMRI (19,29). The parahippocampal region (paraHPC) was included as part of the medial temporal circuitry associated with cortical DMN systems (30–32), whereas the dorsal raphe (DR) was included separately as a serotonergic neuromodulatory region relevant to classical psychedelic action (19). For each rat and session, mean regional BOLD time series were extracted and pairwise Pearson correlations generated 12 × 12 FC matrices. Baseline was compared separately with LSD 15 min and LSD 30 min using paired two-tailed t tests (p < 0.05), with FDR correction across connections (33). Anatomical definitions and ROI abbreviations are provided in Supplementary Table 5.

### Dynamic FC (DFC) analysis

Dynamic FC was estimated across the 12 predefined ROIs using overlapping Gaussian-tapered sliding windows (445 s; 2-s step [1 TR]), yielding 223 windows per session. Within each window, pairwise Pearson correlations were used to construct weighted FC matrices, from which nodal strength (sum of edge weights) and eigenvector centrality (neighbor-weighted nodal influence) were derived (34,35). Negative correlations were set to zero before eigenvector-centrality estimation, whereas strength was computed separately for positive and negative weights. Metric time series were normalized to the corresponding session mean. Baseline was compared separately with LSD 15 min and LSD 30 min at each window using paired t tests (p < 0.05), with Benjamini– Hochberg FDR correction within each ROI’s window series; ROIs were ranked by the number of FDR-significant windows. Complementary session-wise deviations exceeding |z| > 3 were counted descriptively and were not considered statistically significant (34,35).

### Dynamic Causal Modeling (DCM) analysis

Spectral DCM assessed effective connectivity among six predefined ROIs—ACC, PL, IL, HPC, paraHPC, and RSC—at baseline and 15 and 30 min after LSD administration (36,37). For each rat, a fully connected model was estimated in SPM12 from cross-spectral densities, and A-matrix parameters were entered into group-level analyses. Within-session connectivity was assessed using one-sample t tests with FDR correction (q < 0.05). Drug effects were tested separately for baseline versus LSD 15 min and baseline versus LSD 30 min using paired t tests with FDR correction across directed connections. A complementary Bayesian analysis estimated non-zero drug effects (PP > 0.95). The 15 largest connection changes were ranked descriptively.

## Results

### Data-driven ICA reveals distinct LSD-associated network profiles across post-injection periods

Data-driven ICA provided an initial brain-wide view of LSD-associated network reorganization, identifying 100 stable components spanning cortical and subcortical systems (Fig. 2A–B). Component-to-component FC showed widespread LSD-associated changes at both post-injection periods (Fig. 2C). At 15 min, significant changes were biased toward increases (148 increases, 113 decreases), with ACC/RSP the highest-ranked altered component and PFC/NAc, amygdala, hypothalamic, and cerebellar components also prominent (Fig. 2D). At 30 min, the corresponding baseline contrast showed a relative shift toward decreases (111 increases, 151 decreases), accompanied by greater prominence of hippocampal-related components; PAG/PRN/MRN, ACC/RSP, somatosensory, and Hipp/Hab components were also highly ranked. Thus, relative to their respective baseline contrasts, the early profile showed an increase-biased pattern with ACC/RSP prominence, whereas the later profile showed more connectivity decreases and stronger hippocampal-centered network prominence.

**Figure 2.**
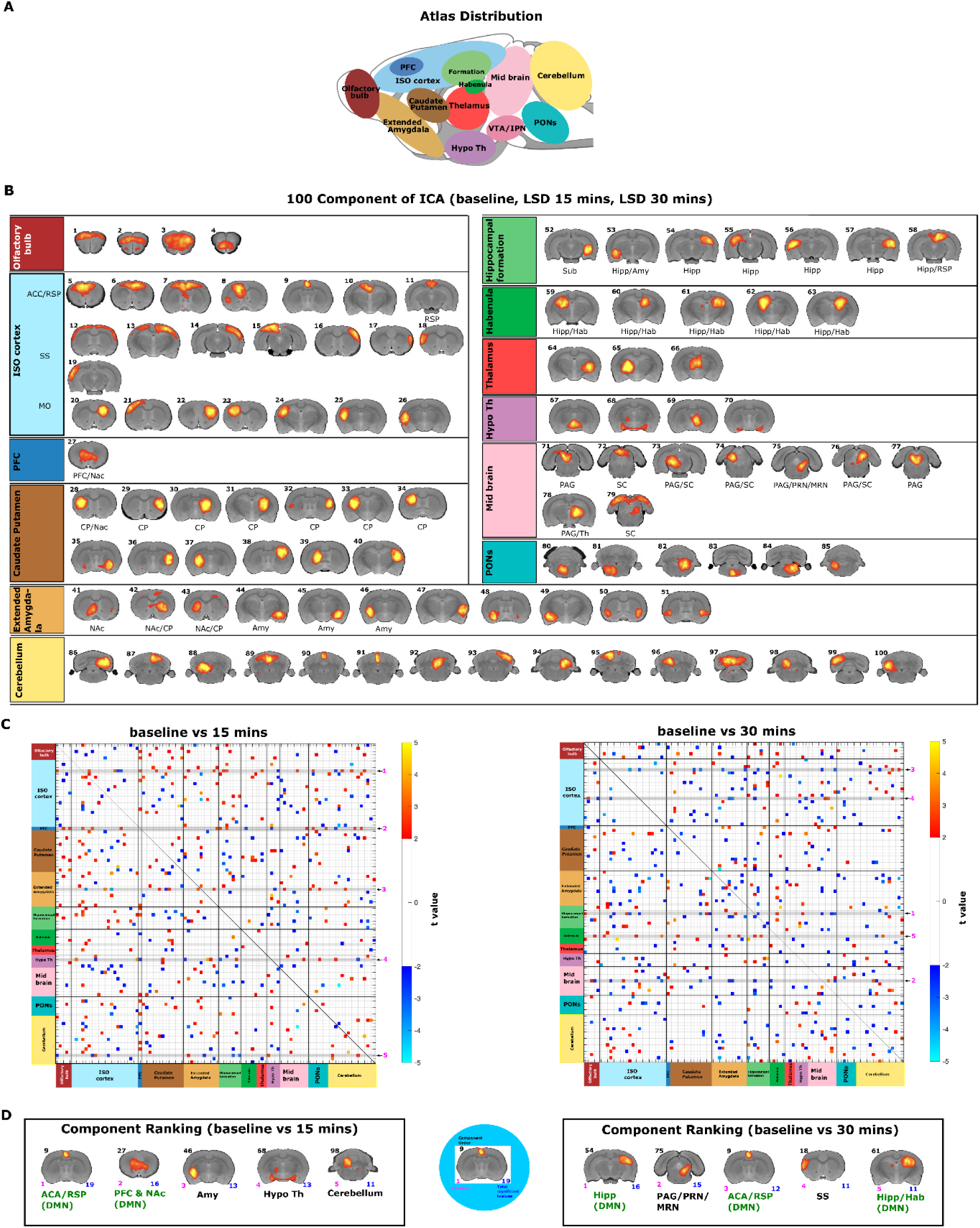
Whole-brain ICA-based functional connectivity across baseline and post-LSD periods. (A) Schematic sagittal representation of the rat brain showing the anatomical distribution of the 13 color-coded macro-regions used to classify the ICA components. (B) Spatial ICA performed on the pooled dataset comprising baseline, 15-min post-LSD, and 30-min post-LSD sessions identified 100 stable components that were subsequently anatomically annotated. Components are displayed in rostrocaudal order and grouped according to their anatomical correspondence with the macro-regions shown in panel A. (C) Whole-brain component-to-component functional connectivity differences relative to baseline. Matrices show significant functional connectivity differences for baseline versus 15 min post-LSD (left) and baseline versus 30 min post-LSD (right). Colors represent t values, with blue/cyan indicating decreased functional connectivity and red/yellow indicating increased functional connectivity relative to baseline. The numbers shown to the right of each matrix indicate a descriptive component ranking based on the total number of significant altered connections involving each component. (D) Top-ranked ICA components according to the number of significant altered functional connections at 15 min post-LSD (left) and 30 min post-LSD (right) relative to baseline. For each component, the black number indicates the ICA component number, the magenta number its rank, and the blue number the total number of significant altered functional connections. The highlighted component illustrates its anatomical localization within the full ICA component distribution. Abbreviations: ACC, anterior cingulate cortex; Amy, amygdala; CP, caudate putamen; ExA, extended amygdala; FC, functional connectivity; Hab, habenula; Hipp, hippocampus; HPF, hippocampal formation; HyTh, hypothalamus; ICA, independent component analysis; LSD, lysergic acid diethylamide; MO, medial orbital cortex; NAc, nucleus accumbens; PAG, periaqueductal gray; PRN, Pontine Reticular Nucleus; MRN, Median Raphe Nucleus; PFC, prefrontal cortex; RSP, retrosplenial cortex; SC, superior colliculus; SS, somatosensory cortex; Th, thalamus.

### Seed-to-voxel FC reveals spatially distributed and seed-specific LSD-associated changes

To complement this data-driven overview with seed-specific whole-brain mapping, seed-to-voxel FC was examined using six predefined seeds (Fig. 3A). Relative to baseline, all six seeds showed widespread increases and decreases across cortical, striatal, thalamic, sensory, and limbic territories at 15 min (Fig. 3B–G). In the 30-min baseline contrast, significant effects persisted, although several seed maps appeared spatially more restricted, whereas paraHPC retained widespread clusters (Fig. 3B–G).

**Figure 3.**
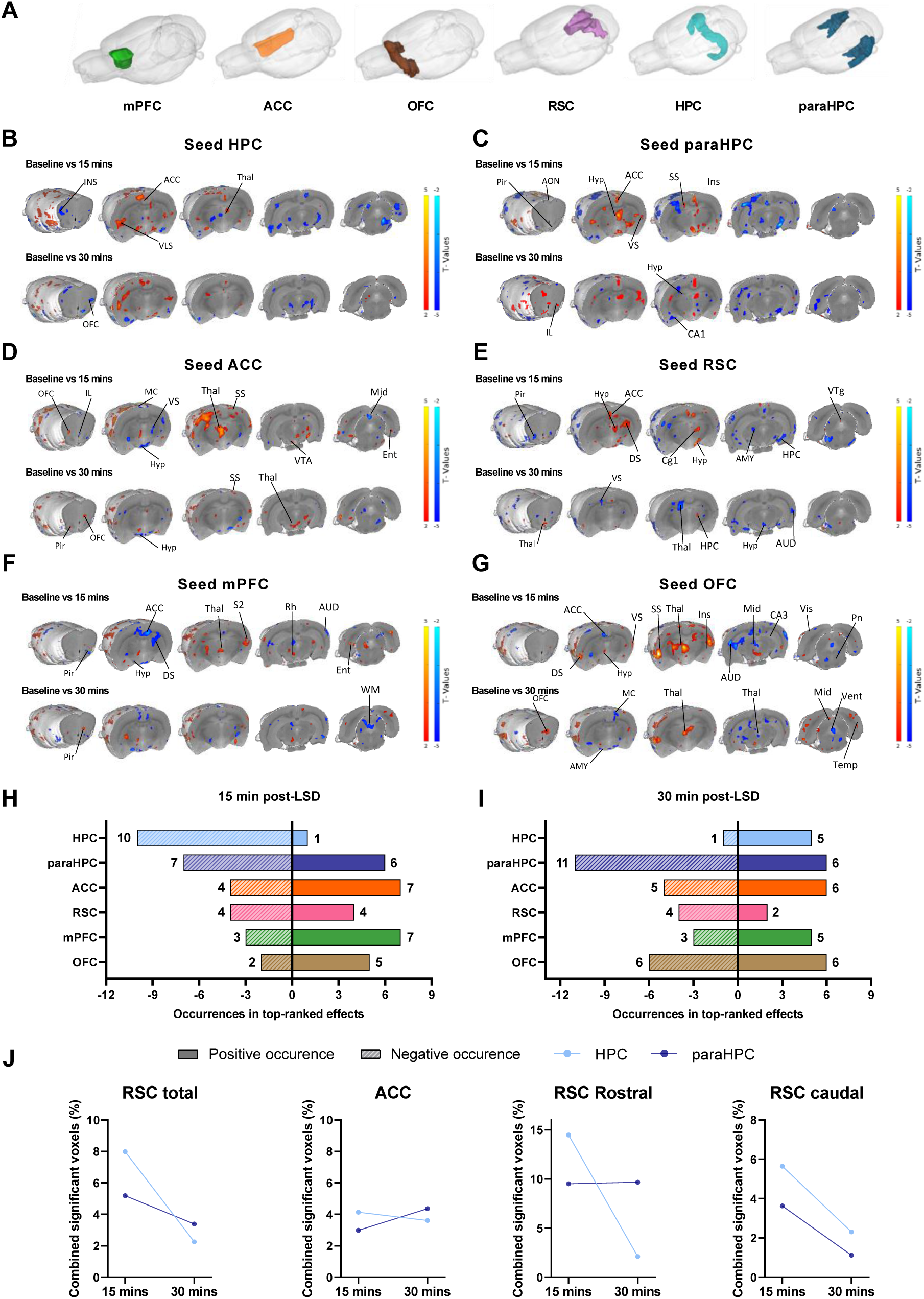
Spatial and quantitative characterization of LSD-associated seed-to-voxel functional connectivity changes across post-injection periods. (A) Anatomical locations of the six DMN-related seeds used for seed-to-voxel analyses: medial prefrontal cortex (mPFC), anterior cingulate cortex (ACC), orbitofrontal cortex (OFC), retrosplenial cortex (RSC), hippocampus (HPC), and parahippocampal region (paraHPC). (B–G) Seed-to-voxel statistical maps showing significant functional connectivity differences relative to baseline for the 15-min and 30-min post-LSD contrasts for the HPC (B), paraHPC (C), ACC (D), RSC (E), mPFC (F), and OFC (G) seeds. Warm-colored clusters indicate increased functional connectivity relative to baseline, whereas cool-colored clusters indicate decreased functional connectivity. Color scales represent t values, and anatomical labels indicate the principal regions encompassed by significant clusters. (H, I) Descriptive counts of negative and positive occurrences among the top-ranked seed-to-voxel seed-target effects for each seed at 15 min (H) and 30 min (I) post-LSD. Solid bars represent positive occurrences and hatched bars represent negative occurrences. (J) Percentage of significant voxels within selected target regions for HPC-and paraHPC-seeded contrasts at 15 and 30 min post-LSD. Quantification is shown for total RSC, total ACC, rostral RSC, and caudal RSC, with light and dark blue lines representing HPC and paraHPC seeds, respectively. Together, the spatial maps and quantitative summaries indicate anatomically heterogeneous seed-specific connectivity changes across the two post-injection baseline contrasts, with prominent hippocampal and parahippocampal involvement.

Quantification of the top-ranked effects revealed distinct seed profiles (Fig. 3H–I). At 15 min, HPC, paraHPC, and ACC were most frequently represented among positive and negative top-ranked effects. At 30 min, paraHPC and OFC were most represented; descriptively, HPC shifted from predominantly negative to predominantly positive representation, while paraHPC remained strongly represented among decreases. For HPC- and paraHPC-seeded maps, the proportion of significant voxels within RSC-related targets was generally lower in the 30-min than in the 15-min baseline contrast, whereas the corresponding proportion within ACC-related targets changed little (Fig. 3J). These analyses quantify the distribution and spatial extent of significant effects, not effect magnitude or formal 15-versus-30-min differences.

### Seed-to-seed stationary FC within DMN-related brain regions

To move from whole-brain seed maps to pairwise interactions within the predefined 12-region network, stationary seed-to-seed FC was examined (Fig. 4A). Global FC distributions are shown across sessions, with the LSD-15 distribution appearing descriptively more compact than baseline and LSD-30 (Fig. 4B). Scatterplots relating baseline and post-LSD connectivity values are shown in Figure 4C. Baseline connectivity involved medial frontal, retrosplenial, temporal association, and HPC/paraHPC nodes (Fig. 4D). At 15 min, the pattern was mixed, with reduced or undetected paraHPC-, RSC-, and temporal/frontal connections alongside strengthened or newly detected sensory-associative links (Fig. 4D–E). At 30 min, the medial frontal core remained detectable, while several paraHPC-centered connections were present, including paraHPC–OFC, paraHPC– AUD, and paraHPC–HPC. VIS-related coupling included VIS–RSC, VIS–paraHPC, and VIS– AUD, and a negative ACC–HPC connection was detected (Fig. 4D–E).

**Figure 4.**
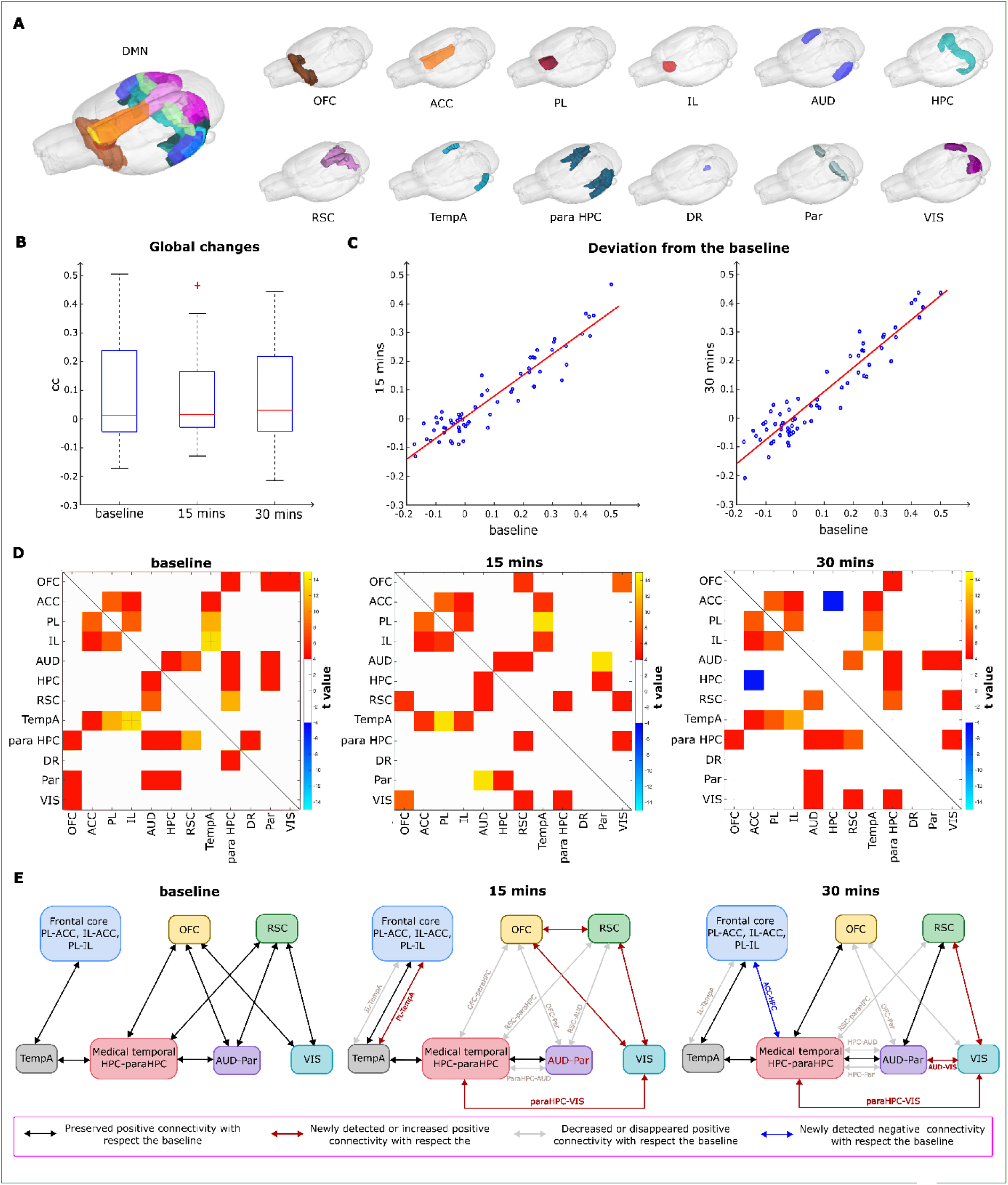
Seed-to-seed stationary functional connectivity within the 12-region DMN-related network across baseline and post-LSD periods. (A) Schematic 3D representation of the 12 DMN-related seed regions mapped onto a rat brain template. A composite view of the full network is shown on the left, and the anatomical location of each individual seed is shown on the right. (B) Boxplots showing the distribution of global seed-to-seed correlation coefficients (cc) across baseline, LSD-15, and LSD-30 sessions. Global FC distributions are shown across sessions, with the LSD-15 distribution appearing descriptively more compact than baseline and LSD-30. (C) Scatter plots illustrating the relationship between baseline connectivity values and post-LSD connectivity values for the LSD-15 (left) and LSD-30 (right) baseline contrasts. Red lines indicate linear regression fits. (D) Within-session seed-to-seed functional connectivity matrices for baseline (left), LSD-15 (middle), and LSD-30 (right). Heatmaps display significant pairwise functional connectivity within the 12-region DMN-related network, with color indicating the corresponding t values for each connection. (E) Schematic summary of the reorganization of seed-to-seed stationary functional connectivity after LSD administration. The diagrams summarize a preserved frontal core across sessions, a 15-min profile characterized by reduced paraHPC- and selected RSC-related coupling together with increased or newly detected sensory-associative interactions, and a 30-min profile in which several paraHPC-centered connections were present, visual-related coupling remained represented, and a negative ACC–HPC connection was detected. Arrow colors indicate preserved positive connectivity (black), newly detected or increased positive connectivity relative to baseline (red), decreased or lost positive connectivity relative to baseline (gray), and newly detected negative connectivity relative to baseline (blue). Abbreviations: OFC, orbitofrontal cortex; ACC, anterior cingulate cortex; PL, prelimbic cortex; IL, infralimbic cortex; AUD, auditory cortex; HPC, hippocampus; RSC, retrosplenial cortex; TempA, temporal association cortex; paraHPC, parahippocampal region; DR, dorsal raphe; Par, parietal association cortex; VIS, visual cortex.

### Dynamic FC reveals distinct medial frontal–hippocampal profiles across post-injection periods

Because stationary FC averages connectivity across each run, dynamic FC was used to resolve time-varying, region- and metric-specific deviations from baseline (Fig. 5A–C). FDR-significant windows were distributed throughout the first post-LSD run, whereas node-strength changes in the 30-min baseline contrast were more concentrated early in the acquisition (Fig. 5A). At 15 min, node strength was most frequently altered in HPC (16.8% of windows), ACC (13.9%), and RSC (9.1%), whereas eigenvector centrality was most frequently altered in ACC (13.0%), PL (8.7%), and RSC (6.7%) (Fig. 5A–B). This profile emphasized hippocampal changes in overall coupling and medial frontal changes in relative network influence. In the 30-min baseline contrast, strength alterations were greatest in IL (8.7%), AUD (6.7%), and HPC (5.3%), while centrality alterations were greatest in HPC (10.6%) and IL (9.1%); ACC showed no FDR-significant centrality windows. Regional rankings and representative strength trajectories are shown in Figure 5B–C. Descriptively, the later baseline contrast was characterized by greater hippocampal and infralimbic centrality involvement.

**Figure 5.**
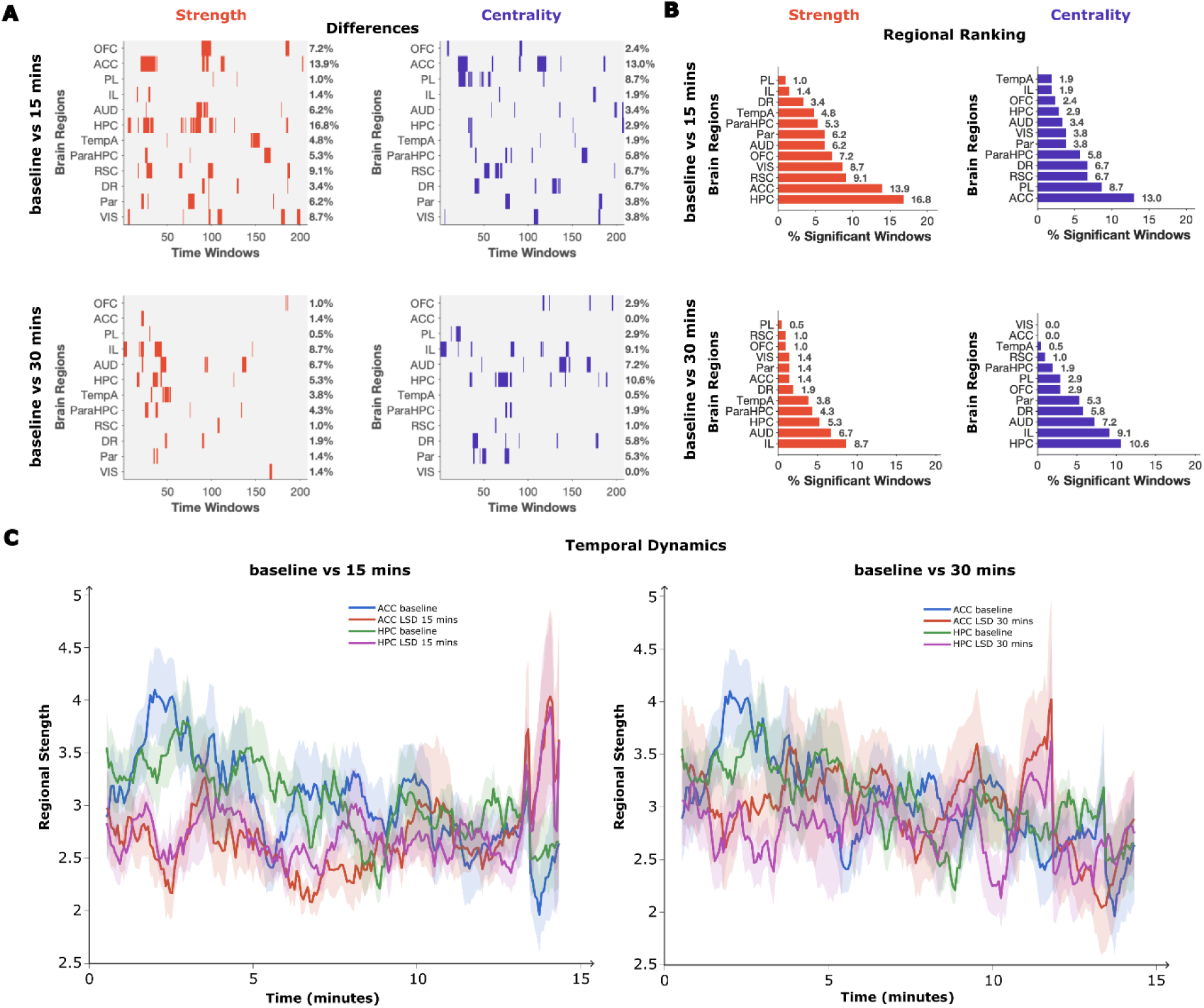
Dynamic functional connectivity alterations within DMN-related regions across post-LSD periods. (A) Binary significance matrices showing time-resolved dynamic functional connectivity differences relative to baseline. For the baseline versus LSD-15 and baseline versus LSD-30 contrasts, node-strength alterations are shown in red and eigenvector-centrality alterations in blue. Rows correspond to the 12 predefined ROIs and columns to successive sliding windows. Colored marks indicate windows showing FDR-significant differences from baseline for the corresponding metric. Percentages indicate the proportion of FDR-significant windows for each region. (B) Descriptive regional ranking of dynamic functional connectivity alterations. Bar plots show the percentage of FDR-significant windows for node strength (red) and eigenvector centrality (blue) for the baseline versus LSD-15 and baseline versus LSD-30 contrasts. Regions are ranked according to the proportion of FDR-significant windows for each metric. (C) Temporal profiles of node strength in representative regions selected from the regional ranking. Time courses show mean normalized regional node strength across the scanning session for baseline and LSD-15 (left) and baseline and LSD-30 (right). Shaded areas indicate SEM across animals. Abbreviations: OFC, orbitofrontal cortex; ACC, anterior cingulate cortex; PL, prelimbic cortex; IL, infralimbic cortex; AUD, auditory cortex; HPC, hippocampus; TempA, temporal association cortex; paraHPC, parahippocampal region; RSC, retrosplenial cortex; DR, dorsal raphe; Par, parietal association cortex; VIS, visual cortex; DFC, dynamic functional connectivity; LSD, lysergic acid diethylamide.

### DCM reveals time-dependent reconfiguration of directional effective connectivity within DMN-related regions under LSD

Because functional connectivity does not establish directionality, DCM was used to characterize directed interactions within the six-region DMN-related circuit (PL, ACC, IL, RSC, HPC, and paraHPC; Fig. 6A). At 15 min, the highest-ranked parameter changes were predominantly decreases in effective connectivity (Fig. 6B–C, upper panels; Supplementary Figure 2). RSC→IL showed the largest reduction and was the only pathway surviving FDR correction, providing the strongest evidence for an early attenuation of posterior-to-frontal influence. Other highly ranked decreases involved RSC→ACC, paraHPC→HPC, and hippocampal outputs to ACC, IL, and PL, whereas increases were less prominent and included PL→ACC, ACC→IL, and paraHPC→PL. Thus, the early DCM profile was descriptively dominated by reduced directed influences, with inferential support strongest for RSC→IL.

**Figure 6.**
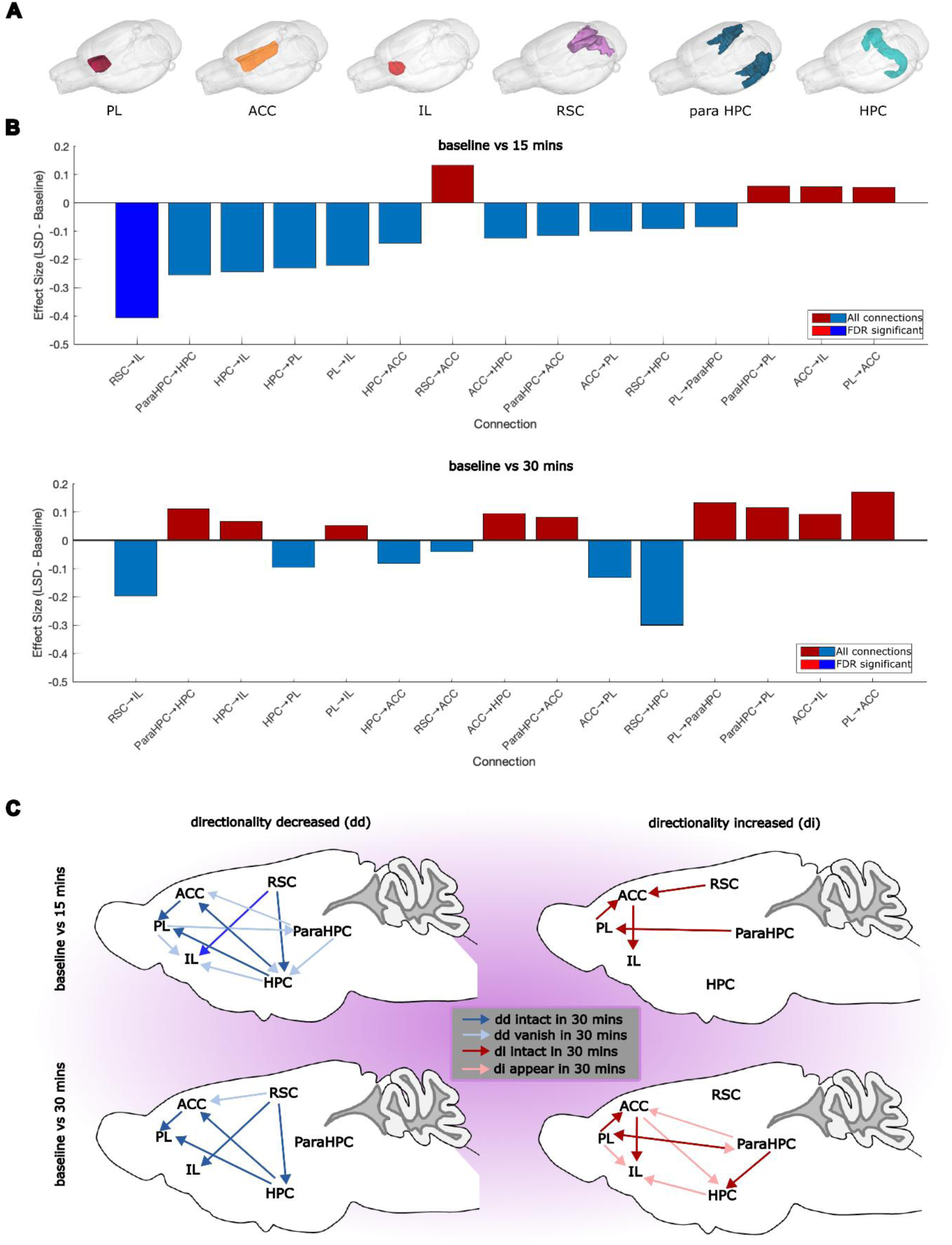
LSD-induced reorganization of directional effective connectivity within the DMN-related network assessed by dynamic causal modeling (DCM). (A) Schematic 3D representation of the six DMN-related regions included in the DCM analysis, mapped onto the rat brain template: prelimbic cortex (PL), anterior cingulate cortex (ACC), infralimbic cortex (IL), retrosplenial cortex (RSC), parahippocampal region (paraHPC), and hippocampus (HPC). (B) Bar plots showing the top 15 directed connections ranked according to the magnitude of the LSD − baseline parameter change for the baseline versus 15-min post-LSD contrast (upper panel) and baseline versus 30-min post-LSD contrast (lower panel). Positive values (red) indicate increased effective connectivity relative to baseline, whereas negative values (blue) indicate decreased effective connectivity. Brighter bars denote connections surviving FDR correction; the RSC→IL decrease at 15 min was the only pathway surviving FDR correction. (C) Schematic summary of LSD-associated reorganization of directional effective connectivity. Left panels illustrate decreases in directed effective connectivity and right panels illustrate increases for the 15-min (upper row) and 30-min (lower row) baseline contrasts. At 15 min, the directional profile was predominantly characterized by decreased effective connectivity, whereas the 30-min profile showed a more balanced distribution of increases and decreases, with greater involvement of hippocampal and parahippocampal pathways. Dark blue arrows indicate decreases represented at both post-LSD periods, whereas light blue arrows indicate decreases represented predominantly in the 15-min profile. Dark red arrows indicate increases represented at both post-LSD periods, whereas light red arrows indicate increases represented predominantly in the 30-min profile. Together, these analyses reveal a marked reorganization of directional interactions within the DMN-related circuit following acute LSD administration, characterized by an early predominance of reduced directed influences and a later profile with more selective hippocampal–parahippocampal and medial frontal reorganization. Abbreviations: ACC, anterior cingulate cortex; DCM, dynamic causal modeling; DMN, default mode network; HPC, hippocampus; IL, infralimbic cortex; LSD, lysergic acid diethylamide; paraHPC, parahippocampal region; PL, prelimbic cortex; RSC, retrosplenial cortex.

In the 30-min baseline contrast, the highest-ranked changes showed a more balanced distribution of increases and decreases (Fig. 6B–C, lower panels; Supplementary Figure 2). Prominent decreases included RSC→HPC and ACC→PL, whereas increases involved HPC→RSC, paraHPC→HPC, and ACC→HPC. The opposing RSC→HPC decrease and HPC→RSC increase descriptively indicated asymmetric reweighting of this pathway. Overall, the 30-min profile was consistent with more selective hippocampal, parahippocampal, and medial frontal reorganization; these pathway-level patterns remained exploratory because they did not survive correction for multiple comparisons.

Figure 7 summarizes the complementary network profiles captured across ICA, seed-to-voxel, stationary FC, dynamic FC, and DCM for the two baseline contrasts.

**Figure 7.**
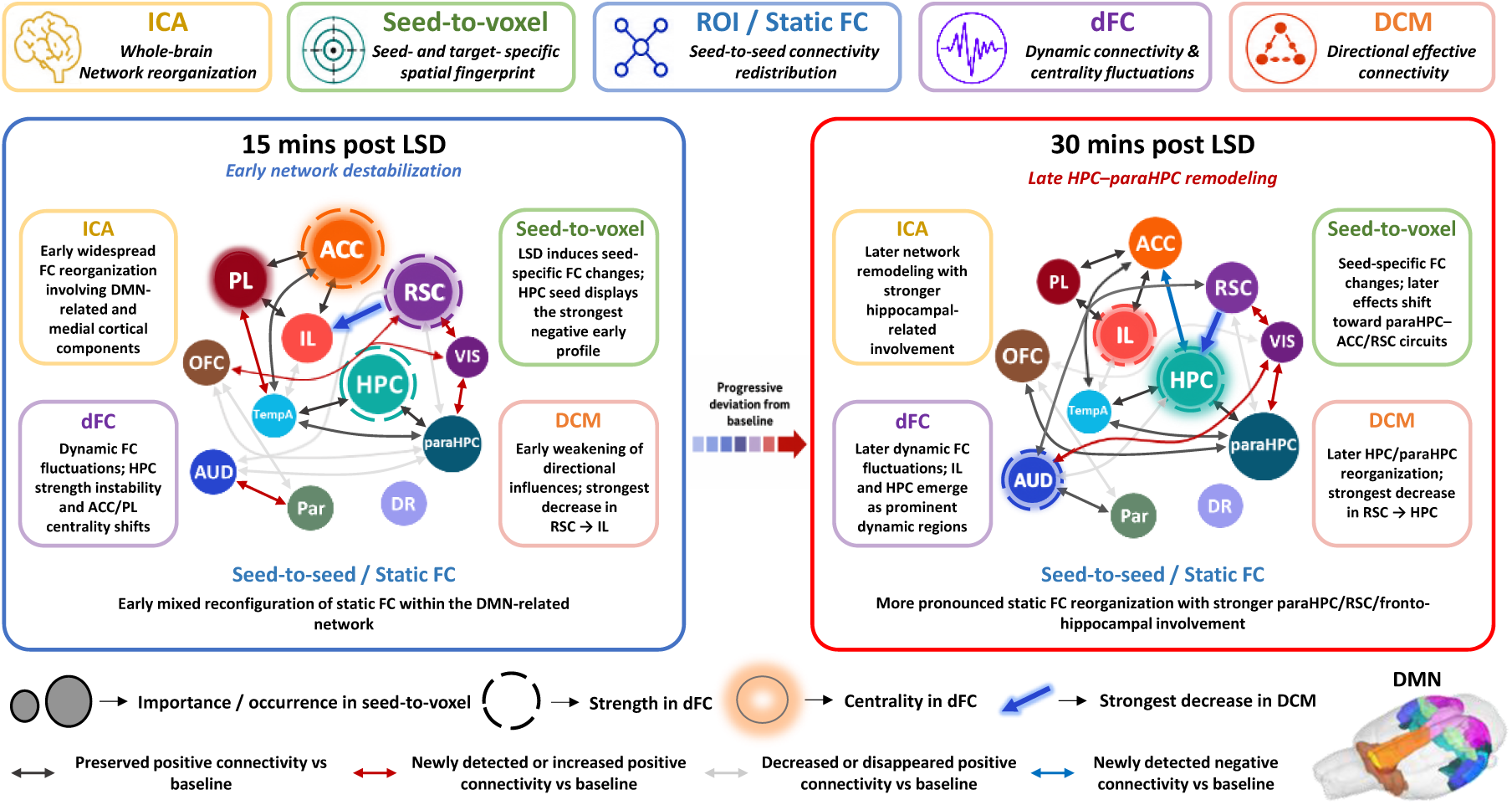
Multimodal summary of DMN-related network reorganization across post-LSD periods. Schematic integration of DMN-related network changes identified in the baseline versus 15-min and baseline versus 30-min post-LSD contrasts across ICA, seed-to-voxel FC, stationary seed-to-seed FC, dynamic FC (dFC), and DCM analyses. Node size reflects regional prominence in ICA and seed-to-voxel analyses, colored highlighting indicates dFC eigenvector centrality, and dashed contours indicate dFC node strength. The blue arrow denotes the strongest FDR-corrected decrease identified by DCM (RSC→IL), whereas the remaining arrows summarize seed-to-seed stationary FC patterns relative to baseline. Overall, the multimodal profile supports a relative shift from broader early network disruption toward greater later hippocampal–parahippocampal involvement. Abbreviations: ACC, anterior cingulate cortex; AUD, auditory cortex; DCM, dynamic causal modeling; dFC, dynamic functional connectivity; DMN, default mode network; DR, dorsal raphe; FC, functional connectivity; HPC, hippocampus; ICA, independent component analysis; IL, infralimbic cortex; LSD, lysergic acid diethylamide; OFC, orbitofrontal cortex; Par, parietal association cortex; paraHPC, parahippocampal region; PL, prelimbic cortex; ROI, region of interest; RSC, retrosplenial cortex; TempA, temporal association cortex; VIS, visual cortex.

## Discussion

A recent systematic review concluded that classical psychedelics consistently alter DMN connectivity, although the mechanistic significance of these changes remains incompletely understood (6). Our findings place these DMN alterations within broader large-scale network reorganization extending beyond canonical DMN boundaries. Here, combining ICA, seed-based functional connectivity, dynamic graph-theoretical measures, and DCM across the initial 30-min post-injection period, we show the onset and early evolution of temporally structured network reorganization. Across analyses, the two baseline contrasts suggested a relative shift from widespread cortical and medial frontal perturbations earlier toward greater hippocampal– parahippocampal involvement later. Effective connectivity further revealed altered directed interactions within the medial cortical–hippocampal network. Together, these findings provide evidence that acute LSD induces a temporally organized reconfiguration of large-scale brain networks, accompanied by a redistribution of directed influences within medial cortical– hippocampal circuitry and a relative shift toward greater hippocampal control rather than uniform disruption of functional connectivity (Fig. 7).

Our findings indicate that psychedelic-induced network reorganization unfolds as a structured temporal process rather than a stationary state, with consecutive post-injection periods showing distinct profiles in the balance of connectivity increases and decreases, affected regions, temporal stability, and direction of interregional influences. To our knowledge, this is the first study combining temporally resolved functional and effective connectivity analyses to characterize successive phases of acute psychedelic network reorganization.

This temporal interpretation is consistent with previous work showing that psychedelics expand the repertoire of functional brain states (16), reorganize connectome-harmonic dynamics (38), exert time-specific effects on dynamic integration and segregation (12), and reduce control energy needed for transitions between recurrent brain states (39). Within this temporally sensitive framework, reduced within-network coherence, particularly within the DMN (6,9), can be reconciled with increased between-network coupling or global functional connectivity (5,11,40). Rather than mutually exclusive, these observations may reflect different analytical dimensions, acquisition windows, or phases of an evolving network response. Our findings therefore underscore the importance of extended, temporally resolved acquisitions for psychedelic DMN studies, because successive configurations may be obscured by time-averaged analyses (12), while effective connectivity can help resolve their directional reorganization. Because the primary inferential contrasts compared each post-LSD period with baseline rather than directly with one another, early–later differences should be interpreted descriptively.

A particularly consistent feature across analyses was a shift in the anatomical center of gravity of the LSD response from widespread cortical perturbation toward greater hippocampal– parahippocampal involvement during the later post-injection window. This is consistent with the medial temporal subsystem’s role in integrating contextual and mnemonic information with cortical DMN hubs (30–32), and with human evidence that LSD alters parahippocampal–retrosplenial connectivity (10). The later medial temporal prominence may therefore reflect relative reweighting of communication between contextual-memory systems and medial cortical DMN-related nodes. This shift suggests that hippocampal circuits may assume greater control over large-scale network organization as the acute psychedelic state evolves. In the absence of behavioral correlates, its functional and experiential consequences cannot be determined directly.

Dynamic FC further showed that LSD-associated alterations were temporally structured rather than uniformly sustained. During the first post-LSD run, significant windows were distributed across the acquisition, whereas in the second run node-strength alterations were concentrated earlier, with centrality changes also occurring later, descriptively suggesting stronger dynamic perturbation in the earlier period. Early changes emphasized HPC coupling strength and ACC/PL centrality, whereas the later profile showed greater HPC/IL centrality involvement, with additional sensory, parietal, and brainstem contributions. Together, these profiles suggest evolving regional network prominence, with medial frontal involvement earlier and greater hippocampal–infralimbic prominence later. Nevertheless, the sliding-window approach limits precise inference about onset and peak timing (41). This temporally structured pattern is consistent with time-specific psychedelic effects on network integration and reduced constraints on transitions between functional configurations (12,39), although comparison with human findings is limited by species and analytical differences and by anesthesia effects on resting-state FC (42).

DCM provided a key mechanistic perspective on network reorganization, moving beyond descriptive FC to characterize how directed influences are redistributed following LSD administration (13,15). At 15 min, the reduction in RSC→IL influence survived FDR correction, providing the strongest evidence for an early alteration in directed posterior-to-frontal communication. Additional highly ranked parameter changes suggested a broader pattern of reduced posterior- and hippocampal-to-frontal coupling, including RSC→ACC and hippocampal outputs toward medial frontal regions, although these effects did not survive correction and remain descriptive. Later parameter estimates suggested selective hippocampal reweighting of directed interactions toward medial temporal circuitry, consistent with a shift toward greater hippocampal control within the later network profile; these later effects also remained below corrected significance.

A potential mechanistic interpretation is that the observed pattern partly reflects 5-HT2A-mediated modulation of cortical neuronal gain and local excitation–inhibition balance. Ketanserin blocks major LSD-induced connectivity alterations, whose spatial distribution corresponds to cortical HTR2A expression (3). Transcriptomics-informed modelling likewise suggests that 5-HT2A-related pyramidal gain can reproduce the regional topography of LSD-induced FC changes (43), while whole-brain effective-connectivity analyses reveal altered interregional coupling and self-inhibition under LSD (15). Such gain modulation may destabilize established interactions and reduce constraints separating functional configurations. Because receptor occupancy and pharmacological blockade were not assessed and LSD acts at additional receptor targets (4), this remains a testable mechanistic hypothesis rather than a demonstration of 5-HT2A dependence. Our DCM findings are compatible with the REBUS framework, as the shift toward greater hippocampal influence may reflect relaxed hierarchical constraints on medial temporal circuitry (44). This shift could involve greater hippocampal contribution to contextual and mnemonic processing, allowing internally generated or memory-related information to exert more influence on large-scale network organization during the acute psychedelic state. Future studies combining rs-fMRI with behavioral measures will be important to clarify the functional significance of this reorganization.

The present findings suggest that the DMN should not be considered the exclusive substrate of psychedelic action but rather one component of a broader hierarchical network reorganization. Instead, LSD appears to selectively reorganize interactions within and beyond the network, consistent with the established fractionation of the DMN into subsystems with distinct anatomical and functional properties (30,31,45). ICA and seed-to-voxel mapping further revealed striatal, thalamic, limbic, and sensory-associative involvement, indicating that DMN-related reorganization was embedded within broader cortico-striato-thalamo-cortical and limbic circuitry. This is consistent with human studies showing that LSD redistributes connectivity between associative and sensory systems and alters thalamic and striatal interactions with posterior cingulate and temporal regions (3,11,14). Together, these findings support large-scale reorganization involving DMN hubs, hippocampal circuitry, and subcortical and sensory systems rather than isolated DMN disruption.

Several limitations should be considered. This exploratory repeated-measures pharmaco-fMRI study used a within-subjects design in males only, with all analyses derived from the same dataset and no time-matched vehicle group. This framework requires cautious cross-method interpretation. Seed-based and DCM analyses were restricted to predefined DMN-related regions, enabling targeted investigation of medial cortical–hippocampal interactions while potentially overlooking effects outside the selected circuit and retaining dependence on ROI definition. Dynamic FC relied on sliding windows, whose results depend on window definition and statistical testing procedures (41,46). Finally, the study examined only the acute post-injection period and lacked behavioral correlates, limiting interpretation of longer-term and functional consequences.

Anesthesia is an inherent consideration in preclinical rs-fMRI because isoflurane can modulate spontaneous BOLD signals, and functional connectivity (47,48), including DMN organization (49). Nevertheless, BOLD signals and resting-state network organization remain detectable under isoflurane across rodent fMRI studies (27,28,48,50,51). Anesthesia-related variability was minimized by maintaining isoflurane at 1.5–1.8% during functional acquisitions, continuously monitoring physiological parameters, and using each animal as its own baseline control. Alternative anesthetic protocols, including dex/medetomidine-based regimens, can also differentially affect functional connectivity (47,52–54), whereas awake imaging may introduce stress and head-motion confounds (42,55). Thus, anesthesia-related effects cannot be excluded, but standardized anesthetic conditions and the repeated-measures design reduce their potential contribution to the observed post-LSD differences.

The later medial temporal prominence may therefore reflect relative reweighting of communication between contextual-memory systems and medial cortical DMN-related nodes. This is particularly relevant to recent human evidence linking psychedelic-induced integration of internally and externally directed processing to self- and boundary-dissolving experiences, conceptualized as embeddedness, together with context-dependent reconfiguration of hippocampal–DMN effective connectivity (56). Our findings therefore raise the possibility that hippocampal reweighting contributes to the altered integration of contextual and internally generated information during the psychedelic state.

Taken together, our findings support a unified mechanistic model of acute LSD action in which large-scale brain reorganization unfolds across successive phases. The early phase appears to correspond to a transient relaxation of directed cortical interactions, whereas the later profile suggests progressive hippocampal-centered reorganization of effective network architecture. Importantly, increases in FC and reductions in effective connectivity describe different properties of network organization and are therefore not directly comparable. FC reflects the strength of statistical coupling between regional signals, whereas DCM estimates the direction and strength of influences between regions. Their combination therefore provides complementary information on both the organization and directional reconfiguration of the network over time. One apparent divergence in the psychedelic neuroimaging literature is that reduced within-network coherence, particularly within the DMN (6,10), can coexist with increased between-network or global connectivity (5,40).

Rather than simply disrupting the DMN, acute LSD induces a temporally organized reconfiguration of large-scale brain hierarchy characterized by a shift toward hippocampal-centered control of network organization. By integrating functional and effective connectivity across successive phases of the acute psychedelic state, the present study provides a mechanistic framework that may help explain how psychedelic-induced network reorganization emerges over time.

## Supporting information

Supplementary Material, Methods and Results

## Acknowledgements

We would like to thank Prof. Luc Mallet and the ADELY consortium for their contribution to the Treatment of Alcohol DEpendence with LYsergic acid diethylamide: translational approach and clinical efficacy study (ADELY) project.

This research was funded by IReSP and INCa as part of the 2021 call for research projects aimed at combating the use of and addiction to psychoactive substances. Project numbers: SPAV1-22-019 and IRESP-AAPSPA2021-V1-04. INSERM (French National Institute of Health and Medical Research) and Hauts-de-France Regional Council doctoral fellowship to FFH.

FFH: Conceptualization, Methodology, Investigation, Formal analysis, Visualization, writing – original draft, Writing – review & editing, Project administration. RB: Methodology, Formal analysis, Data curation, Software, Visualization. JJ: Conceptualization, Methodology, Resources, Funding acquisition, Writing – review & editing. MS: Methodology, Formal analysis, Data curation, Software, Writing – review & editing. SF: Investigation, Writing – original draft. RU, PK, SS: Formal analysis, Writing – review & editing. MN: Conceptualization, Methodology, Resources, Writing – original draft, Writing – review & editing, Supervision, Project administration, Funding acquisition. SBH: Conceptualization, Methodology, Resources, Writing – original draft, Writing – review & editing, Supervision, Project administration. MTN: Conceptualization, Methodology, Formal analysis, Data curation, Software, Visualization, Validation, Writing – original draft, Writing – review & editing, Supervision, Project administration.

The datasets generated and analyzed during the current study, as well as the custom analysis scripts, are available from the corresponding authors on reasonable request, pending approval from the institutional ethics committee and in compliance with the INSERM data-sharing policies.

This study was performed in compliance with EU Directive 2010/63/EU and French Decree No. 2013-118, and received ethical approval from the local animal ethics committee (CREMEAP; APAFIS #26615). All experiments adhered to the ARRIVE guidelines and the 3Rs principle. Animals were housed under standard conditions with ad libitum food/water, and isoflurane anesthesia with continuous physiological monitoring was applied to minimize stress and ensure animal welfare throughout the MRI acquisitions.

