## Supplementary Material, Methods and Results for "Acute lysergic acid diethylamide induces a time-dependent shift toward hippocampal control of default mode network reorganization"

### Supplementary Methods and Materials

#### Animals and housing

Animals were maintained under a 12-hour light/dark cycle (lights on at 7:00 AM) at 22–23°C and 45–55% humidity. Rats were housed individually in a cage including a wooden chew stick and a cardboard tube. Rats were handled for 5 days after arrival following the 3H guidelines (https://www.3hs-initiative.co.uk/the-3hs/handling), and MRI experiments began 5 days later. Animals were naïve before the study. MRI experiments were performed at the small-animal MRI facility (PIRMPA), Université de Picardie Jules Verne, Amiens, France. No expected or unexpected adverse events were observed.

#### Image acquisition

Animals were initially anesthetized with isoflurane (5% in air; 1 L/min) in an induction chamber for up to 2 min before being placed in prone position on a Bruker rat bed. Isoflurane anesthesia was used to minimize motion and acute restraint-related stress and to maintain stable physiological conditions during continuous baseline and post-injection acquisitions. The head was stabilized using a bite bar and ear bars, and anesthesia was delivered through a face mask.

Body temperature (36.86 ± 0.22°C) and respiration were monitored continuously using an MR-compatible monitoring system (SA Instruments, Stony Brook, NY, USA). Body temperature was maintained within physiological limits using a warm-water circulation system (Bruker, Ettlingen, Germany). Oxygen saturation and heart rate (382 ± 8.85 bpm) were recorded using a fiber-optic pulse oximeter attached to the left hind paw. Vaporizer output was finely adjusted to stabilize respiratory rate at approximately 80 ± 3 breaths/min. Physiological parameters are reported in Supplementary Figure 1.

MRI was performed on a BioSpec 70/20 USR 7 T system (Bruker, Ettlingen, Germany) operating at 300 MHz with AV-III electronics and ParaVision 360 v3.4. A four-channel phased-array receive-only surface coil was combined with a 72-mm transmit-only volume resonator, and magnetic field homogeneity was optimized using FASTMAP shimming. An 8-min anatomical scan preceded the functional acquisitions.

Functional BOLD images were acquired in the coronal plane using an SE-EPI sequence with the following parameters: TR = 2000 ms, TE = 16 ms, flip angle = 90°, 450 repetitions, FOV = 30 × 25 $mm^2$, matrix = 64 × 64, 34 slices, in-plane spatial resolution = 0.48 x 0.40 $mm^2$ and slice thickness = 0.8 mm. The Bruker standard EPI navigator was used with automatic ghost correction and drift compensation.

Following the 15-min baseline BOLD acquisition, the sequence was paused for 90 s to allow remote intraperitoneal LSD injection through a 70-cm polyethylene tube connected to a 23 G (0.6 × 25 mm) needle, enabling administration without disturbing the scanner bed. Isoflurane was briefly increased to 2% immediately before injection to minimize the risk of transient arousal and associated motion following LSD administration. It was then lowered to 1.5–1.8% one minute before acquisition resumed. Two consecutive 15-min post-injection BOLD runs were then acquired. Successful injection was confirmed at the end of each session by verifying needle position and the absence of subcutaneous liquid accumulation.

Anatomical $T_2$-weighted images were obtained using a fast spin-echo (TurboRARE) sequence (RARE factor = 8; TR/TE = 2740/44 ms; FOV = 35 × 30 $mm^2$; matrix = 256 × 218; 28 contiguous coronal slices of 0.8mm of thickness; in-plane spatial resolution = 0.14 x 0.14 $mm^2$, NEX = 4). Total acquisition time per animal, including baseline, post-drug, and anatomical scans, was approximately 55 min.

### Image preprocessing

Raw Bruker fMRI data were converted to NIfTI format before preprocessing. Skull stripping was performed using the Smart Swin Transformer combined with a Dense U-Net architecture (1). Functional images were subsequently denoised using a 3D U-WGAN (Wasserstein Generative Adversarial Network with a 3D Dense U-Net discriminator) (2).

The first five volumes of each dataset were discarded to allow for magnetic field stabilization. Spike removal was performed using AFNI's 3dDespike (3,4) with local editing parameters, replacing outlier intensities with the mean of adjacent non-outlier values. Motion correction was then performed using AFNI's 3dvolreg, with the first volume as reference, and the resulting motion parameters were retained for subsequent nuisance regression. Slice-timing correction was applied using FSL 6.0.6 (5) to adjust for inter-slice acquisition delays based on an interleaved acquisition scheme. Bias field correction was implemented using ANTs' N4BiasFieldCorrection to correct for low-frequency intensity inhomogeneities.

For anatomical registration and segmentation, each rat's fMRI dataset was aligned to an in-house built 3D MRI Rat Brain Atlas (© Ekam Solutions, Boston, MA, USA), which comprises 174 segmented and annotated brain regions, using ANTs' registration tools. Nuisance regression was performed using FSL and AFNI and included motion parameters together with mean time series extracted from ventricular and white matter masks (1,6,7). Spatial smoothing was performed using AFNI's blurmaster_input approach with a Gaussian kernel (FWHM = 0.4 mm). Finally, temporal filtering was performed with AFNI using a band-pass filter (0.01–0.1 Hz) to isolate low-frequency fluctuations relevant to neuronal activity (6–14).

### Postprocessing: Independent Component Analysis

Independent component analysis was performed using GIFT (v1.3i) implemented in MATLAB. Data from all nine rats and all three sessions—baseline, LSD 15 min, and LSD 30 min—were analyzed as a single group dataset, following previously established procedures (15–17). A 100-component ICA decomposition was applied to identify LSD- and DMN-related anatomical components. Component stability and reliability were assessed using ICASSO with 20 repetitions based on randomization and bootstrapping (16).

Following component estimation and spatial sorting, FC analysis was conducted to examine differences between baseline and LSD conditions (15 min and 30 min) across anatomically annotated components, in accordance with our previous methodologies (1,15–17). MATLAB was used for time-series extraction, FC computation, and statistical testing. The time series corresponding to each of the 100 components from the three sessions were used to generate component-to-component FC matrices. Pearson correlation coefficients were computed for each component pair, converted to Fisher Z-scores, and tested for significance using one-sample t-tests (threshold $t > 4$; $p < 0.01$; $n = 9$). Comparisons between baseline and LSD conditions (15 min and 30 min) were conducted using pairwise two-tailed t-tests (threshold $t > 2$; $p = 0.05$). False discovery rate (FDR) correction was applied to control for multiple comparisons. Finally, to evaluate the relative importance of each component, a component-ranking analysis was performed by counting the total number of significant FC differences involving that component (16,17).

Formal tests of distributional assumptions were not performed across individual imaging outcomes. Where applicable, correlation coefficients were Fisher Z-transformed before group-level parametric testing.

### Postprocessing: Seed-to-seed stationary FC analysis

The selected ROIs encompassed medial prefrontal, retrosplenial, hippocampal/parahippocampal, sensory-associative, and serotonergic neuromodulatory regions relevant to DMN-like organization and psychedelic action. Bilateral or midline seed masks were derived from anatomical masks based on the 3D Rat Brain Atlas© (Ekam Solutions, Boston, MA), comprising 174 segmented and annotated brain regions (1,6,7). When necessary, adjacent anatomical subregions were combined using FSL tools to generate the final ROI masks.

For each animal and each session, corresponding to baseline, 15 min post-LSD, and 30 min post-LSD, the mean BOLD time series was extracted from each ROI using an in-house MATLAB pipeline. Pairwise Pearson correlation coefficients were then computed between all ROI time series to generate 12 × 12 stationary FC matrices for each session. Statistical testing of pairwise connections was performed using two-tailed t-tests, and significance was assessed using a threshold of $p < 0.05$ with FDR correction. Significant positive and negative connections were retained for matrix visualization and network-level interpretation. Scatterplots with regression lines, histograms, and boxplots were generated using MATLAB built-in functions to visualize changes in correlation structure, distributions of connectivity values, and session-dependent differences.

### Postprocessing: Dynamic FC (DFC) analysis

Dynamic FC was estimated separately for each rat and session (baseline, LSD 15 min, and LSD 30 min) across the 12 predefined DMN-related ROIs using an overlapping Gaussian-tapered boxcar window of 445 s and a 2-s step (1 TR), yielding 223 windows per session. Within each window, pairwise Pearson correlations between ROI time series were computed to generate a weighted FC matrix. Two nodal graph-theoretical measures were derived for each window: strength, defined as the sum of a node's edge weights, and eigenvector centrality, which weights a node's influence according to the centrality of its neighbors (13,14). As eigenvector centrality requires a non-negative adjacency matrix, negative correlation weights were set to zero prior to centrality estimation; node strength was computed separately for positive and negative weights to preserve information from anti-correlated connections. Dynamic FC analyses were performed using an in-house MATLAB pipeline.

For each ROI, group-level (across-animal) strength and centrality time series were normalized to their respective baseline or post-LSD session mean. Baseline-versus-LSD differences were assessed at each time window using paired (within-subject) t-tests ($\alpha = 0.05$), reflecting the repeated-measures design in which each animal contributed both baseline and post-injection data; this was performed independently for the 15-min and 30-min sessions. Because sliding windows overlap and adjacent time points are therefore not statistically independent, and because tests were performed across 12 ROIs and 223 windows per session, a Benjamini–Hochberg false discovery rate correction was applied within each ROI's window series prior to ranking. The number of significant windows per ROI was summed to rank ROIs by their sensitivity to LSD-induced connectivity changes.

As a complementary descriptive analysis, window-wise values were z-scored within each session using the corresponding session mean and standard deviation. Windows with $|z| > 3$ were classified as substantial deviations, and their number was used as an additional descriptive ranking measure; these windows were not considered statistically significant.

### Postprocessing: Dynamic Causal Modeling (DCM) analysis

Spectral DCM was performed in SPM12 implemented in MATLAB, following our previous study (8), to estimate directed effective connectivity among the ACC, PL, IL, HPC, paraHPC, and RSC at baseline and 15 and 30 min after LSD administration (18,19). The Statistical Parametric Mapping (SPM) tool (https://www.fil.ion.ucl.ac.uk/spm/software/spm12/) was used in MATLAB to calculate DCM following our previous study (8). A fully connected model was specified for each rat (n = 9) and inverted using spm_dcm_fmri_csd, which estimates effective connectivity by modeling observed cross-spectral densities. Subject-level parameter estimates from the A matrices were then extracted for group-level analysis.

For each session, one-sample t-tests were performed on each connection, with significance determined using FDR correction ($q < 0.05$). LSD-induced changes were directly assessed using paired t-tests comparing baseline with each post-LSD session for each directed connection, with FDR correction applied across the tested connections. In addition, a complementary Bayesian analysis was performed to estimate the posterior probability of a non-zero drug effect for each connection, using PP > 0.95 as the threshold. Significant effects were classified as increased or decreased under LSD relative to baseline.

Results were visualized using group-mean connectivity matrices, t-statistic maps, and directed network graphs. As a complementary descriptive summary, the 15 connections showing the largest effect sizes were ranked and reported together with their direction of change and statistical significance.

### Postprocessing: Seed-to-voxel analysis

Seed-to-voxel correlation analyses were performed using the FSL toolbox. For each subject (rat) and session, the mean BOLD time series was extracted from each predefined seed region and correlated voxel-wise with the time series of all other brain voxels to generate whole-brain FC maps. The resulting correlation coefficients were converted to Fisher's Z scores to improve normality.

Group-level statistical analyses were conducted using one-sample t-tests (threshold: $t > 4$, $p < 0.01$, n = 9) to identify regions showing significant positive connectivity. Baseline-versus-LSD within-subject comparisons were performed using pairwise t-tests (threshold: $t > 2$, $p < 0.05$, one-tailed) to assess connectivity differences between baseline and LSD-treated groups at 15- and 30-min post-administration. All group-level statistical maps were corrected for multiple comparisons using a family-wise error rate (FWER) correction applied at the cluster level ($p < 0.05$). Visualization of FC maps was performed using BrainNet Viewer (https://www.nitrc.org/projects/bnv/) and Mango (http://ric.uthscsa.edu/mango/) for 3D rendering and surface projection onto a standardized rat brain template.

For quantitative analyses of seed-to-voxel connectivity within selected brain regions, an in-house MATLAB script (12,16) was employed to extract mean Z-values within anatomical ROIs. The resulting data were visualized using BrainNet Viewer for 3D representation overlaid on the 3D Rat Brain Atlas©.

#### Seed-to-voxel quantification and ranking workflow

Seed-to-voxel effects were quantified from the statistical maps generated for each seed region and the two post-LSD time points. Six seeds were considered: ACC, HPC, OFC, paraHPC, mPFC, and RSC. For each seed, 174 anatomical target masks were evaluated separately at 15- and 30-min post-LSD relative to baseline.

For each seed-target pair, the number of significant positive voxels was defined as voxels with $t >= +2.0$, whereas the number of significant negative voxels was defined as voxels with $t <= -2.0$. The combined number of significant voxels corresponded to all voxels with $|t| >= 2.0$. Positive, negative, and combined percentages were computed by dividing the corresponding voxel counts by the total number of voxels within each anatomical target mask and multiplying by 100. Thus, all percentages represent the spatial extent of significant modulation within a target region rather than subject-level effect sizes.

To reduce the full anatomical list to a concise set of the most prominent effects, seed-target pairs were ranked separately for each time point and direction of effect. At each time point, positive effects were ranked according to the percentage of positive significant voxels, and negative effects were ranked according to the percentage of negative significant voxels. The top 30 seed-target pairs were retained for each of the four summaries: positive effects at 15 min (Supplementary Table 1), negative effects at 15 min (Supplementary Table 2), positive effects at 30 min (Supplementary Table 3), and negative effects at 30 min (Supplementary Table 4).

The ranking-based bar graph was generated by counting, within each top-30 list, how many times each seed appeared among the strongest positive or negative effects. This analysis was used as a descriptive index of seed representation among the most prominent seed-to-voxel changes and was not used as an inferential statistical test (Figure 3H-I).

HPC and paraHPC were selected for target-specific follow-up analyses because the ranking-based summary suggested a temporal redistribution between these medial temporal seeds. ACC and RSC were selected as target regions because of their relevance to rat DMN-like circuitry and because they provided clear examples of time-dependent changes in HPC- and paraHPC-associated connectivity (Figure 3J).

### Supplementary figures and tables

A

| Parameters | Baseline | 5–15 min | 20–30 min |
|---|---|---|---|
| Temperature, °C | 36.07 ± 0.22 | 37.19 ± 0.32 | 37.50 ± 0.18 |
| Respiration, /min | 72.89± 5.24 | 83.39 ± 8.80 | 87.85 ± 5.17 |
| Heart rate | 409.37 ± 11.9 | 359.83 ± 15.5 | 376.31 ± 9.46 |

***Supplementary Figure 1. Physiological monitoring during baseline and post-LSD acquisitions.*** *Group summary of body temperature, respiratory rate, and heart rate during baseline and the 5–15 min and 20–30 min post-injection periods. Values are presented as mean ± SEM (n = 9).*

| 15 min Positive | | | | | | | | | | | |
|---|---|---|---|---|---|---|---|---|---|---|---|
| Rank | Timepoint | Seed | Target region | Ranking metric (%) | Positive voxels | Negative voxels | Combined voxels | Total voxels in mask | Positive % | Negative % | Combined % |
| 1 | 15 min | OFC | central medial thalamic nucleus | 60.37 | 291 | 0 | 291 | 482 | 60.37 | 0 | 60.37 |
| 2 | 15 min | paraHPC | paraventricular nucleus | 50.32 | 310 | 0 | 310 | 616 | 50.32 | 0 | 50.32 |
| 3 | 15 min | OFC | reuniens nucleus | 49.07 | 422 | 0 | 422 | 860 | 49.07 | 0 | 49.07 |
| 4 | 15 min | ACC | central medial thalamic nucleus | 48.55 | 234 | 0 | 234 | 482 | 48.55 | 0 | 48.55 |
| 5 | 15 min | paraHPC | dorsal medial nucleus | 45.86 | 133 | 0 | 133 | 290 | 45.86 | 0 | 45.86 |
| 6 | 15 min | mPFC | central medial thalamic nucleus | 45.44 | 219 | 0 | 219 | 482 | 45.44 | 0 | 45.44 |
| 7 | 15 min | mPFC | reuniens nucleus | 41.98 | 361 | 0 | 361 | 860 | 41.98 | 0 | 41.98 |
| 8 | 15 min | ACC | red nucleus | 36.59 | 161 | 0 | 161 | 440 | 36.59 | 0 | 36.59 |
| 9 | 15 min | OFC | medial dorsal thalamic nucleus | 36.36 | 336 | 0 | 336 | 924 | 36.36 | 0 | 36.36 |
| 10 | 15 min | OFC | ventrolateral thalamic nucleus | 35.74 | 614 | 0 | 614 | 1718 | 35.74 | 0 | 35.74 |
| 11 | 15 min | paraHPC | suprachiasmatic nucleus | 35.71 | 30 | 0 | 30 | 84 | 35.71 | 0 | 35.71 |
| 12 | 15 min | paraHPC | ventral medial nucleus | 35 | 210 | 0 | 210 | 600 | 35 | 0 | 35 |
| 13 | 15 min | paraHPC | extended amygdala | 33.7 | 246 | 0 | 246 | 730 | 33.7 | 0 | 33.7 |
| 14 | 15 min | ACC | medial dorsal thalamic nucleus | 30.63 | 283 | 0 | 283 | 924 | 30.63 | 0 | 30.63 |
| 15 | 15 min | mPFC | posterior hypothalamic area | 29.93 | 261 | 0 | 261 | 872 | 29.93 | 0 | 29.93 |
| 16 | 15 min | ACC | posterior hypothalamic area | 29.01 | 253 | 0 | 253 | 872 | 29.01 | 0 | 29.01 |
| 17 | 15 min | ACC | primary somatosensory ctx shoulde | 27.82 | 276 | 0 | 276 | 992 | 27.82 | 0 | 27.82 |
| 18 | 15 min | mPFC | ventral tegmental area | 27.12 | 166 | 0 | 166 | 612 | 27.12 | 0 | 27.12 |
| 19 | 15 min | RSC | ventral anterior thalamic nucleus | 26.63 | 220 | 0 | 220 | 826 | 26.63 | 0 | 26.63 |
| 20 | 15 min | RSC | retrochiasmatic nucleus | 26.19 | 55 | 0 | 55 | 210 | 26.19 | 0 | 26.19 |
| 21 | 15 min | RSC | paraventricular nucleus | 25.97 | 160 | 0 | 160 | 616 | 25.97 | 0 | 25.97 |
| 22 | 15 min | mPFC | supramammillary nucleus | 25.77 | 101 | 0 | 101 | 392 | 25.77 | 0 | 25.77 |
| 23 | 15 min | mPFC | medial dorsal thalamic nucleus | 24.46 | 226 | 0 | 226 | 924 | 24.46 | 0 | 24.46 |
| 24 | 15 min | OFC | secondary somatosensory ctx | 22.97 | 1314 | 0 | 1314 | 5720 | 22.97 | 0 | 22.97 |
| 25 | 15 min | ACC | reuniens nucleus | 22.79 | 196 | 0 | 196 | 860 | 22.79 | 0 | 22.79 |
| 26 | 15 min | HPC | medial dorsal thalamic nucleus | 22.4 | 207 | 30 | 237 | 924 | 22.4 | 3.25 | 25.65 |
| 27 | 15 min | ACC | anterior thalamic nuclei | 21.13 | 347 | 6 | 353 | 1642 | 21.13 | 0.37 | 21.5 |
| 28 | 15 min | mPFC | red nucleus | 20.91 | 92 | 0 | 92 | 440 | 20.91 | 0 | 20.91 |
| 29 | 15 min | RSC | substantia innominata | 20.87 | 48 | 0 | 48 | 230 | 20.87 | 0 | 20.87 |
| 30 | 15 min | paraHPC | central medial thalamic nucleus | 20.54 | 99 | 0 | 99 | 482 | 20.54 | 0 | 20.54 |

***Supplementary Table 1. Top-ranked positive seed-to-voxel effects at 15 min post-LSD.***

| 15 min Negative | | | | | | | | | | | |
|---|---|---|---|---|---|---|---|---|---|---|---|
| Rank | Timepoint | Seed | Target region | Ranking metric (%) | Positive voxels | Negative voxels | Combined voxels | Total voxels in mask | Positive % | Negative % | Combined % |
| 1 | 15 min | ACC | suprachiasmatic nucleus | 54.76 | 0 | 46 | 46 | 84 | 0 | 54.76 | 54.76 |
| 2 | 15 min | HPC | dorsal raphe | 40.8 | 0 | 142 | 142 | 348 | 0 | 40.8 | 40.8 |
| 3 | 15 min | mPFC | retrochiasmatic nucleus | 40 | 0 | 84 | 84 | 210 | 0 | 40 | 40 |
| 4 | 15 min | ACC | retrochiasmatic nucleus | 37.62 | 0 | 79 | 79 | 210 | 0 | 37.62 | 37.62 |
| 5 | 15 min | ACC | supraoptic nucleus | 32.61 | 1 | 90 | 91 | 276 | 0.36 | 32.61 | 32.97 |
| 6 | 15 min | mPFC | suprachiasmatic nucleus | 32.14 | 0 | 27 | 27 | 84 | 0 | 32.14 | 32.14 |
| 7 | 15 min | RSC | posterior hypothalamic area | 26.38 | 0 | 230 | 230 | 872 | 0 | 26.38 | 26.38 |
| 8 | 15 min | HPC | subthalamic nucleus | 26.1 | 0 | 83 | 83 | 318 | 0 | 26.1 | 26.1 |
| 9 | 15 min | HPC | pontine reticular nucleus oral | 22.01 | 0 | 496 | 496 | 2254 | 0 | 22.01 | 22.01 |
| 10 | 15 min | paraHPC | substantia nigra reticularis | 21.84 | 0 | 359 | 359 | 1644 | 0 | 21.84 | 21.84 |
| 11 | 15 min | HPC | parafascicular thalamic nucleus | 21.11 | 0 | 388 | 388 | 1838 | 0 | 21.11 | 21.11 |
| 12 | 15 min | HPC | ventral posterolateral thalamic nucleus | 21.06 | 2 | 326 | 328 | 1548 | 0.13 | 21.06 | 21.19 |
| 13 | 15 min | paraHPC | medial geniculate | 20.86 | 0 | 237 | 237 | 1136 | 0 | 20.86 | 20.86 |
| 14 | 15 min | paraHPC | supramammillary nucleus | 19.64 | 0 | 77 | 77 | 392 | 0 | 19.64 | 19.64 |
| 15 | 15 min | mPFC | copula of the pyramis | 19.44 | 2 | 340 | 342 | 1749 | 0.11 | 19.44 | 19.55 |
| 16 | 15 min | paraHPC | primary somatosensory ctx trunk | 19.12 | 1 | 283 | 284 | 1480 | 0.07 | 19.12 | 19.19 |
| 17 | 15 min | RSC | ventral tegmental area | 19.12 | 0 | 117 | 117 | 612 | 0 | 19.12 | 19.12 |
| 18 | 15 min | HPC | ventrolateral thalamic nucleus | 18.92 | 5 | 325 | 330 | 1718 | 0.29 | 18.92 | 19.21 |
| 19 | 15 min | HPC | supramammillary nucleus | 18.88 | 1 | 74 | 75 | 392 | 0.26 | 18.88 | 19.13 |
| 20 | 15 min | HPC | CA3 hippocampus ventral | 18.76 | 9 | 355 | 364 | 1892 | 0.48 | 18.76 | 19.24 |
| 21 | 15 min | paraHPC | parietal ctx | 18.64 | 1 | 627 | 628 | 3364 | 0.03 | 18.64 | 18.67 |
| 22 | 15 min | RSC | supramammillary nucleus | 18.62 | 0 | 73 | 73 | 392 | 0 | 18.62 | 18.62 |
| 23 | 15 min | HPC | reticulotegmental nucleus | 18.51 | 0 | 67 | 67 | 362 | 0 | 18.51 | 18.51 |
| 24 | 15 min | paraHPC | prerubral field | 18.38 | 0 | 93 | 93 | 506 | 0 | 18.38 | 18.38 |
| 25 | 15 min | paraHPC | posterior hypothalamic area | 17.78 | 0 | 155 | 155 | 872 | 0 | 17.78 | 17.78 |
| 26 | 15 min | RSC | CA1 hippocampus ventral | 17.42 | 0 | 461 | 461 | 2646 | 0 | 17.42 | 17.42 |
| 27 | 15 min | ACC | anterior hypothalamic area | 17.19 | 0 | 207 | 207 | 1204 | 0 | 17.19 | 17.19 |
| 28 | 15 min | OFC | pedunculopontine tegmental area | 16.92 | 0 | 133 | 133 | 786 | 0 | 16.92 | 16.92 |
| 29 | 15 min | OFC | raphe obscurus nucleus | 16.54 | 0 | 44 | 44 | 266 | 0 | 16.54 | 16.54 |
| 30 | 15 min | HPC | ventral posteromedial thalamic nucleus | 15.88 | 0 | 315 | 315 | 1984 | 0 | 15.88 | 15.88 |

***Supplementary Table 2. Top-ranked negative seed-to-voxel effects at 15 min post-LSD.***

| 30 min Positive | | | | | | | | | | |
|---|---|---|---|---|---|---|---|---|---|---|
| Rank | Timepoint | Seed | Target region | Ranking metric (%) | Positive voxels | Negative voxels | Combined voxels | Total voxels in mask | Positive % | Negative % | Combined % |
| 1 | 30 min | OFC | medial dorsal thalamic nucleus | 42.86 | 396 | 0 | 396 | 924 | 42.86 | 0 | 42.86 |
| 2 | 30 min | paraHPC | paraventricular nucleus | 34.25 | 211 | 0 | 211 | 616 | 34.25 | 0 | 34.25 |
| 3 | 30 min | HPC | suprachiasmatic nucleus | 29.76 | 25 | 0 | 25 | 84 | 29.76 | 0 | 29.76 |
| 4 | 30 min | ACC | ventral tegmental area | 29.58 | 181 | 0 | 181 | 612 | 29.58 | 0 | 29.58 |
| 5 | 30 min | OFC | 1st cerebellar lobule | 26.3 | 142 | 0 | 142 | 540 | 26.3 | 0 | 26.3 |
| 6 | 30 min | OFC | central medial thalamic nucleus | 25.1 | 121 | 0 | 121 | 482 | 25.1 | 0 | 25.1 |
| 7 | 30 min | paraHPC | dorsal medial nucleus | 24.14 | 70 | 0 | 70 | 290 | 24.14 | 0 | 24.14 |
| 8 | 30 min | RSC | medial septum | 24.07 | 103 | 0 | 103 | 428 | 24.07 | 0 | 24.07 |
| 9 | 30 min | paraHPC | facial nucleus | 24.01 | 357 | 25 | 382 | 1487 | 24.01 | 1.68 | 25.69 |
| 10 | 30 min | mPFC | ectorhinal ctx | 23.88 | 277 | 0 | 277 | 1160 | 23.88 | 0 | 23.88 |
| 11 | 30 min | paraHPC | habenula nucleus | 22.08 | 189 | 0 | 189 | 856 | 22.08 | 0 | 22.08 |
| 12 | 30 min | mPFC | intercalated amygdaloid nucleus | 20.59 | 49 | 0 | 49 | 238 | 20.59 | 0 | 20.59 |
| 13 | 30 min | mPFC | red nucleus | 19.77 | 87 | 0 | 87 | 440 | 19.77 | 0 | 19.77 |
| 14 | 30 min | HPC | magnocellular preoptic nucleus | 18.88 | 108 | 1 | 109 | 572 | 18.88 | 0.17 | 19.06 |
| 15 | 30 min | mPFC | ventral tegmental area | 18.14 | 111 | 0 | 111 | 612 | 18.14 | 0 | 18.14 |
| 16 | 30 min | HPC | precuniform nucleus | 17.84 | 86 | 0 | 86 | 482 | 17.84 | 0 | 17.84 |
| 17 | 30 min | HPC | raphe obscurus nucleus | 17.67 | 47 | 0 | 47 | 266 | 17.67 | 0 | 17.67 |
| 18 | 30 min | ACC | intercalated amygdaloid nucleus | 17.65 | 42 | 0 | 42 | 238 | 17.65 | 0 | 17.65 |
| 19 | 30 min | ACC | supramammillary nucleus | 17.6 | 69 | 0 | 69 | 392 | 17.6 | 0 | 17.6 |
| 20 | 30 min | paraHPC | medial orbital ctx | 16.39 | 157 | 0 | 157 | 958 | 16.39 | 0 | 16.39 |
| 21 | 30 min | OFC | periolivary nucleus | 16.28 | 197 | 6 | 203 | 1210 | 16.28 | 0.5 | 16.78 |
| 22 | 30 min | ACC | raphe linear | 15.88 | 88 | 0 | 88 | 554 | 15.88 | 0 | 15.88 |
| 23 | 30 min | paraHPC | medial dorsal thalamic nucleus | 15.58 | 144 | 55 | 199 | 924 | 15.58 | 5.95 | 21.54 |
| 24 | 30 min | ACC | medial cerebellar nucleus fastigia | 15.49 | 70 | 0 | 70 | 452 | 15.49 | 0 | 15.49 |
| 25 | 30 min | mPFC | vestibular nucleus | 15.46 | 489 | 0 | 489 | 3164 | 15.46 | 0 | 15.46 |
| 26 | 30 min | ACC | ectorhinal ctx | 15.34 | 178 | 0 | 178 | 1160 | 15.34 | 0 | 15.34 |
| 27 | 30 min | OFC | trapezoid body | 15.01 | 118 | 0 | 118 | 786 | 15.01 | 0 | 15.01 |
| 28 | 30 min | RSC | substantia innominata | 14.78 | 34 | 0 | 34 | 230 | 14.78 | 0 | 14.78 |
| 29 | 30 min | OFC | cochlear nucleus | 14.18 | 221 | 1 | 222 | 1558 | 14.18 | 0.06 | 14.25 |
| 30 | 30 min | HPC | secondary somatosensory ctx | 13.99 | 800 | 12 | 812 | 5720 | 13.99 | 0.21 | 14.2 |

***Supplementary Table 3. Top-ranked positive seed-to-voxel effects at 30 min post-LSD.***

| 30 min Negative | | | | | | | | | | | |
|---|---|---|---|---|---|---|---|---|---|---|---|
| Rank | Timepoint | Seed | Target region | Ranking metric (%) | Positive voxels | Negative voxels | Combined voxels | Total voxels in mask | Positive % | Negative % | Combined % |
| 1 | 30 min | mPFC | suprachiasmatic nucleus | 82.14 | 0 | 69 | 69 | 84 | 0 | 82.14 | 82.14 |
| 2 | 30 min | ACC | suprachiasmatic nucleus | 79.76 | 0 | 67 | 67 | 84 | 0 | 79.76 | 79.76 |
| 3 | 30 min | paraHPC | dorsal raphe | 54.31 | 0 | 189 | 189 | 348 | 0 | 54.31 | 54.31 |
| 4 | 30 min | mPFC | retrochiasmatic nucleus | 32.86 | 0 | 69 | 69 | 210 | 0 | 32.86 | 32.86 |
| 5 | 30 min | RSC | lateral dorsal thalamic nucleus | 27.03 | 0 | 100 | 100 | 370 | 0 | 27.03 | 27.03 |
| 6 | 30 min | ACC | supraoptic nucleus | 25 | 0 | 69 | 69 | 276 | 0 | 25 | 25 |
| 7 | 30 min | mPFC | supraoptic nucleus | 23.91 | 15 | 66 | 81 | 276 | 5.43 | 23.91 | 29.35 |
| 8 | 30 min | paraHPC | prerubral field | 20.55 | 0 | 104 | 104 | 506 | 0 | 20.55 | 20.55 |
| 9 | 30 min | OFC | dentate gyrus dorsal | 20.39 | 4 | 906 | 910 | 4444 | 0.09 | 20.39 | 20.48 |
| 10 | 30 min | paraHPC | inferior olivary complex | 17.99 | 0 | 145 | 145 | 806 | 0 | 17.99 | 17.99 |
| 11 | 30 min | OFC | CA3 dorsal | 17.87 | 1 | 703 | 704 | 3934 | 0.03 | 17.87 | 17.9 |
| 12 | 30 min | ACC | retrochiasmatic nucleus | 17.62 | 0 | 37 | 37 | 210 | 0 | 17.62 | 17.62 |
| 13 | 30 min | ACC | precuniform nucleus | 15.56 | 0 | 75 | 75 | 482 | 0 | 15.56 | 15.56 |
| 14 | 30 min | paraHPC | substantia nigra compacta | 15.47 | 0 | 103 | 103 | 666 | 0 | 15.47 | 15.47 |
| 15 | 30 min | paraHPC | substantia nigra reticularis | 15.27 | 0 | 251 | 251 | 1644 | 0 | 15.27 | 15.27 |
| 16 | 30 min | RSC | raphe linear | 13.9 | 0 | 77 | 77 | 554 | 0 | 13.9 | 13.9 |
| 17 | 30 min | paraHPC | lemniscal nucleus | 13.78 | 0 | 196 | 196 | 1422 | 0 | 13.78 | 13.78 |
| 18 | 30 min | OFC | medial geniculate | 13.64 | 0 | 155 | 155 | 1136 | 0 | 13.64 | 13.64 |
| 19 | 30 min | OFC | dorsomedial tegmental area | 13.6 | 0 | 108 | 108 | 794 | 0 | 13.6 | 13.6 |
| 20 | 30 min | HPC | medial geniculate | 13.56 | 5 | 154 | 159 | 1136 | 0.44 | 13.56 | 14 |
| 21 | 30 min | paraHPC | primary somatosensory ctx trunk | 13.38 | 1 | 198 | 199 | 1480 | 0.07 | 13.38 | 13.45 |
| 22 | 30 min | paraHPC | primary somatosensory ctx shoulde | 13.31 | 0 | 132 | 132 | 992 | 0 | 13.31 | 13.31 |
| 23 | 30 min | RSC | lemniscal nucleus | 13.22 | 4 | 188 | 192 | 1422 | 0.28 | 13.22 | 13.5 |
| 24 | 30 min | OFC | retrochiasmatic nucleus | 12.86 | 0 | 27 | 27 | 210 | 0 | 12.86 | 12.86 |
| 25 | 30 min | paraHPC | reticulotegmental nucleus | 12.43 | 0 | 45 | 45 | 362 | 0 | 12.43 | 12.43 |
| 26 | 30 min | RSC | CA1 hippocampus ventral | 12.17 | 86 | 322 | 408 | 2646 | 3.25 | 12.17 | 15.42 |
| 27 | 30 min | ACC | ventral medial nucleus | 12.17 | 0 | 73 | 73 | 600 | 0 | 12.17 | 12.17 |
| 28 | 30 min | paraHPC | ventral tegmental area | 11.93 | 0 | 73 | 73 | 612 | 0 | 11.93 | 11.93 |
| 29 | 30 min | paraHPC | lateral amygdaloid nucleus | 11.58 | 0 | 94 | 94 | 812 | 0 | 11.58 | 11.58 |
| 30 | 30 min | OFC | periaqueductal gray thalamus | 11.38 | 0 | 522 | 522 | 4586 | 0 | 11.38 | 11.38 |

***Supplementary Table 4. Top-ranked negative seed-to-voxel effects at 30 min post-LSD.***

**A** **Baseline vs 15 mins**

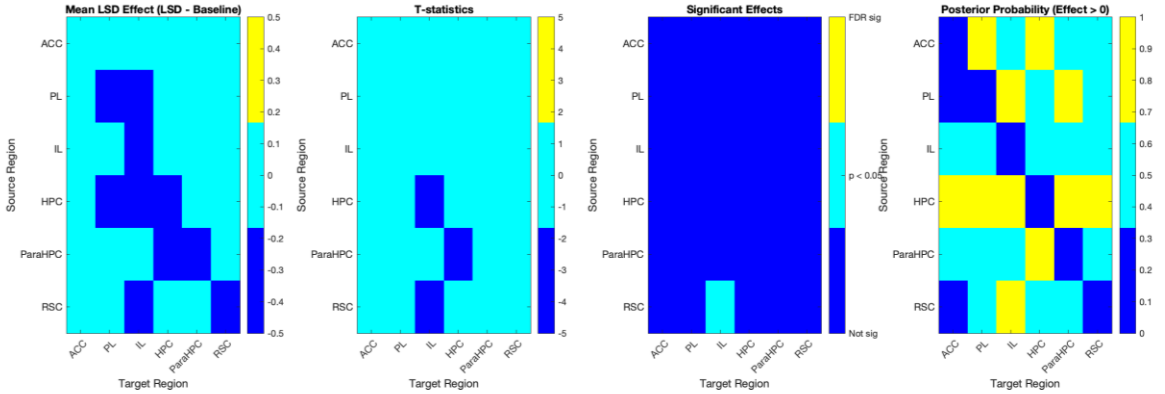


**B** **Baseline vs 30 mins**

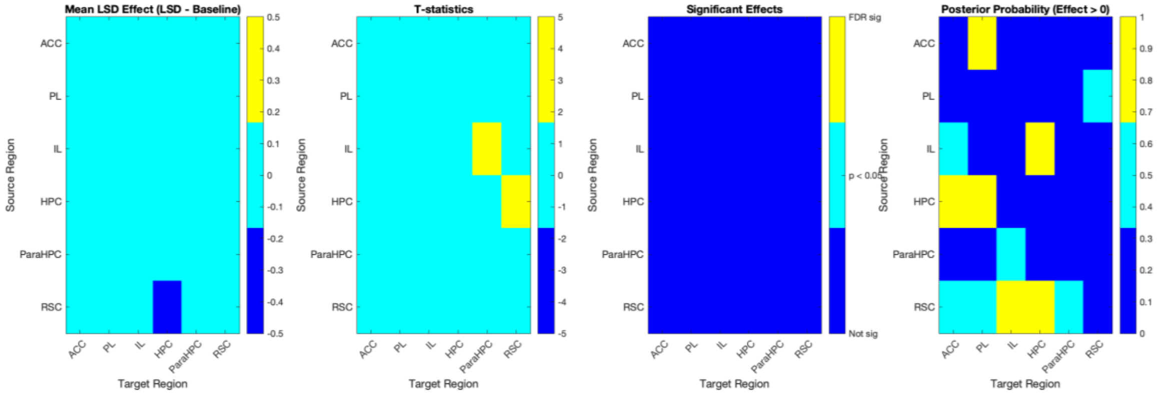


***Supplementary Figure 2. LSD-induced changes in directional effective connectivity within the predefined DCM network.*** *Directional effective connectivity changes are shown for baseline versus 15 min (A) and baseline versus 30 min (B) after LSD administration. For each comparison, matrices show, from left to right, the mean LSD-induced change relative to baseline (LSD − baseline), corresponding t-statistics, FDR-corrected statistical significance, and Bayesian posterior probability that the effect differs from zero. Rows indicate source regions and columns indicate target regions; therefore, each matrix element represents the directed influence from the row region to the column region. Positive values indicate increased and negative values decreased effective connectivity relative to baseline. At 15 min, the RSC→IL decrease was the only connection surviving FDR correction. At 30 min, no connection survived FDR correction, and the observed pathway-level changes should therefore be interpreted as exploratory.*

***Supplementary Table 5. Anatomical identification and structural classification of rat brain regions.***

| Structure image | Structure name | ROI abbreviation | Merged structures |
|---|---|---|---|
| 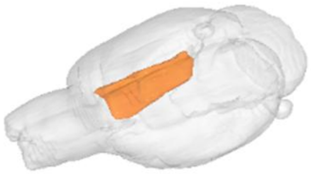 | Anterior cingulate cortex | ACC/ACA | Ø |
| 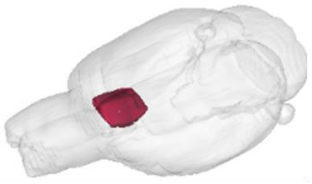 | Prelimbic cortex | PL | Ø |
| 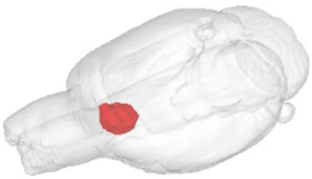 | Infralimbic cortex | IL | Ø |
| 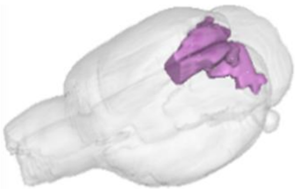 | Retrosplenial cortex | RSC/RSP | Ø |
| 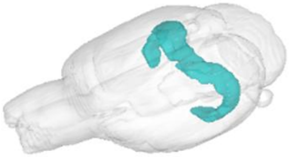 | Hippocampus | HPC | Ø |
| 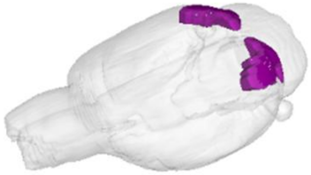 | Visual cortex | VIS | Ø |

| Structure image | Structure name | ROI abbreviation | Merged structures |
|---|---|---|---|
| 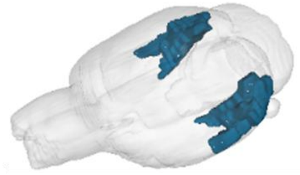 | Parahippocampal cortex | paraHPC | Ø |
| 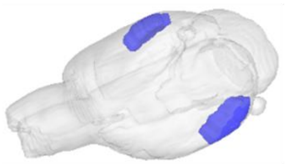 | Auditory cortex | AUD | Ø |
| 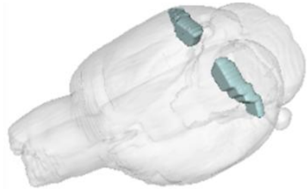 | Parietal cortex | Par | Ø |
| 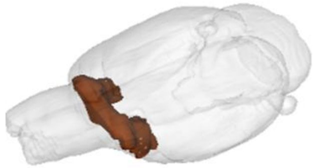 | Orbitofrontal cortex | OFC | Ø |
| 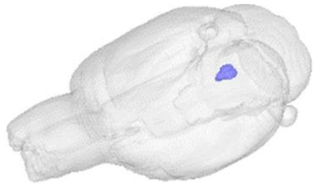 | Dorsal raphe | DR | Ø |
| 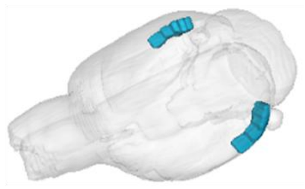 | Temporal association cortex | TempA | Ø |

| Structure image | Structure name | ROI abbreviation | Merged structures |
|---|---|---|---|
| 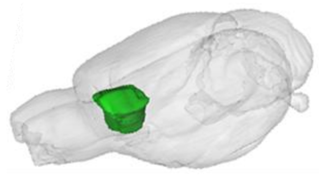 | Medial prefrontal cortex | mPFC | Infralimbic cortex;<br>Prelimbic cortex |
